# Unveiling the Odor Representation in the Inner Brain of *Drosophila* through Compressed Sensing

**DOI:** 10.1101/2023.07.19.549810

**Authors:** Kiri Choi, Won Kyu Kim, Changbong Hyeon

## Abstract

The putative dimension of a space spanned by chemical stimuli is deemed enormous; however, when odorant molecules are bound to a finite number of receptor types and their information is transmitted and projected to a perceptual odor space in the brain, a substantial reduction in dimensionality is made. Compressed sensing (CS) is an algorithm that enables recovery of high-dimensional signals from the data compressed in a lower dimension when the representation of such signals is sufficiently sparse. By analyzing the recent *Drosophila* connectomics data, we find that the *Drosophila* olfactory system effectively meets the prerequisites for CS to work. The neural activity profile of projection neurons (PNs) can be faithfully recovered from a low-dimensional response profile of mushroom body output neurons (MBONs) which can be reconstructed using the electrophysiological recordings to a wide range of odorants. By leveraging the residuals calculated between the measured and the predicted MBON responses, we visualize the perceptual odor space by means of residual spectrum and discuss the differentiability of an odor from others. Our study highlights the sparse coding of odor to the receptor space as an essential component for odor identifiability, clarifying the concentration-dependent odor percept. Further, a simultaneous exposure of the olfactory system to many different odorants saturates the neural activity profile of PNs, significantly degrading the capacity of signal recovery, resulting in a perceptual state analogous to “olfactory white.” Our study applying the CS to the connectomics data provides novel and quantitative insights into the odor representation in the inner brain of *Drosophila*.

## I. INTRODUCTION

Olfaction is a sensory process of detecting chemical stimuli in the environment. Despite its fundamental importance, still left ambiguous are basic quantities associated with the understanding of olfaction, such as the number of differentiable odors and the dimension of odor percepts [1–3]. This is in stark contrast to other sensory processes such as vision, which is quantitatively understood as the combined responses of distinct types of photoreceptors to visual signals continuous in the spectral domain. The odor identification and perception are still deemed highly subjective and context-dependent, varying from one individual to another [4]. However, the conditions for understanding the insect olfactory system have greatly been improved in recent years, thanks to the three-dimensional map of neuronal morphology and the synaptic connectivity reconstructed from high-resolution electron microscopy (EM) images, such as *Drosophila* hemibrain dataset [5], the olfaction-specific FAFB dataset [6], and more recently the full connectome of the *Drosophila* larva [7]. By making the neural circuitry of the insect brain accessible at synapse resolution, these connectomics datasets open a new avenue to a quantitative understanding of the principles behind olfactory processing.

As illustrated in Fig. 1A, olfactory signaling starts when an odorant binds to olfactory receptors (ORs) in the olfactory receptor neurons (ORNs). Each ORN expresses a specific type of OR, exhibiting homotype-dependent responses to input odorants. The same type of ORNs converges to a particular set of uniglomerular PNs (uPNs) forming a glomerulus in the antennal lobe (AL) [8]. The signals converged and sorted at uPNs transfer to the higher olfactory centers such as the mushroom body (MB) calyx and the lateral horn (LH), where synaptic wirings with higher order neurons integrate the signal. Specifically, in the MB calyx, extensive and seemingly randomized signal integration occurs at the synaptic interface between PNs and the Kenyon cells (KCs) [9, 10], which passes over to MBONs whose dimension is smaller than that of PNs. In LH, the compartmentalized spatial organization of PNs translates to a more stereotyped signal integration by lateral horn neurons (LHNs) [11–14}].

**FIG. 1.**
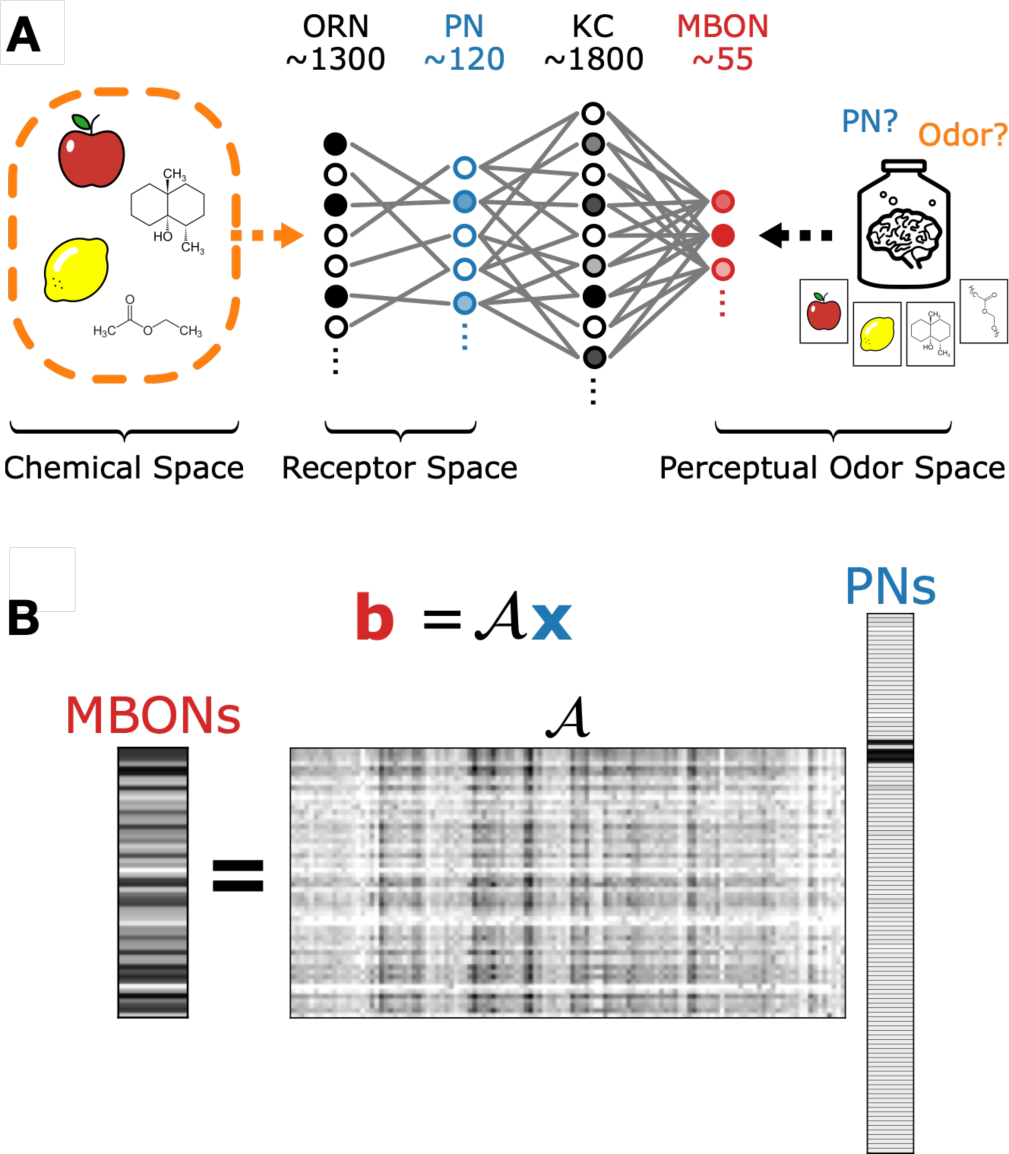
Olfactory processes in *Drosophila*. (A) An odor represented in the chemical space is encoded by a sparse set of ORs/PNs in the receptor space. Highly integrated olfactory signals in the perceptual odor space are deciphered at higher cortical areas, which infers the activity profiles of uPNs (blue) and hence the olfactory code from the relatively low dimensional and randomized information integrated into MBONs (red). (B) Our problem is formulated to solve for *n*-dimensional sparse olfactory codes (**x**) encoded by PNs from a set of underdetermined linear equations in the form of **b** = 𝒜**x**, where **b** and 𝒜 represent the *m*(*< n*)-dimensional response profile of MBONs and the *m* × *n* sensing matrix, respectively.

For a given odor representation at MBONs, there may exist a neural process analogous to solving a constrained optimization problem that enables higher cortical areas to know the identity of the odor in the form of “odor code”, defined as a *combinatorial* expression of uPN activities [10, 15–19] (Fig. 1A). Among several candidate algorithms, compressed sensing (CS), which determines the sparsest solution of a set of underdetermined linear equations, has previously been applied to the *Drosophila* olfactory system as a means to identify the chemical composition of an odor from activity profiles (i.e., neuronal firing rates) of ORNs, PNs, or KCs [17, 20–2 Provided that the chemical composition of an odor represented in the chemical feature space is sparse [24, 25], CS can reliably uncover the odor identity. CS has been applied to the olfactory sensing in the scope of the odorant- OR interface [17, 22, 23] and between odorants and KCs [19, 21]. The previous studies using CS on olfaction have offered valuable insights into the odor codes and odor identification process; yet, they are largely limited to the relationship between the chemical feature space and the receptor space.

In this study, we analyze the neural circuit of the *Drosophila* olfactory system from the connectomics data, and investigate (i) whether the idea of CS [10] enables an accurate reconstruction of odor code from the highly integrated and compressed olfactory information at MBONs (Fig. 1A, B) and (ii) whether the perception of a given odorant(s) is unambiguously discernible from other odors in the perceptual odor space. We test the capacity of CS in recovering uPN activities from MBON response profiles reconstructed from the electrophysiological recordings of neural spiking activities [25, 26]. In particular, we calculate the spectrum of residuals to visualize the perceptual odor space by comparing the reconstructed odor response profiles to those projected onto other olfactory stimuli (see Fig. 2). We explore how the perceptual odor space changes when multiple odorants are mixed, demonstrating that the simultaneous introduction of diverse odorants significantly deteriorates the capacity of CS, an observation in line with the notion of “olfactory white.” We also show that a significant variation in concentration can yield a large deviation in the odor code and alter the extent of odor identifiability. By employing the CS algorithm as a means to probe the olfactory processing in the inner brain of *Drosophila*, our study presents a novel perspective on odor perception while demonstrating the functional benefits of randomized wiring and sparse coding.

**FIG. 2.**
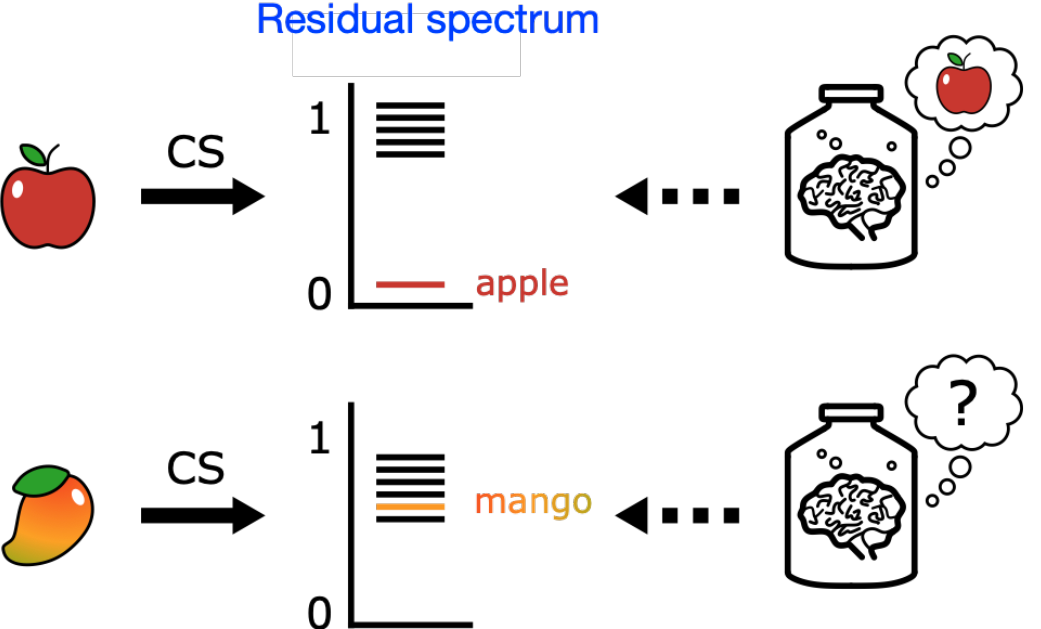
Odor representation and perceptual odor space. A putative structure of the perceptual odor space. The gap between the self-residual and cross-residuals in the residual spectrum decides the differentiability of a given odor (e.g., apple, mango in the illustration) from others.

## II. DATA PREPARATION

The connectomics data analyzed in this study is based on the hemibrain dataset (hemibrain v1.2.1) [5]. When querying the neurons, we chose only PNs, KCs, and MBONs that exhibit at least one synaptic connection. We then collected a total of 119 uPNs synapsing with 1776 KCs, which in turn synapse with 56 MBONs. As for the glomerular labels, we used the one supplied by the hemibrain dataset. In case a uPN was associated with multiple glomeruli, we referred to the labeling from the FAFB dataset [6], which provided their version of the glomerular label, along with the corresponding neuron ID in the hemibrain dataset. Additionally, we updated several labels following the recent community agreement to rename three glomeruli with conflicting glomerulus labels, so as to amend the homotypes VC3l, VC3m, and VC5 to VC3, VC5, and VM6, respectively [27]. In this study, we designate the homotype of a PN by the glomerulus innervated by the dendrites, so that the homotype is uniquely defined for uPNs.

The odor response profile was generated by merging two different datasets that measured the electrophysiological responses to the exposure of an odorant [25, 26]. The experimental procedure used several different concentrations of odorants. 10^−2^ volume per volume (v/v) dilution with either H_2_O, paraffin oil, or mineral oil is used, except for geosmin which used 10^−3^ dilution. We explicitly stated the dilution level whenever we used a different odorant concentration. When there was an overlap between the datasets, we chose the more recent recordings from Seki *et al*. [26] as the representative value. The connection between ORs and the respective glomerulus is identified through a literature search [28– The dataset by Hallem *et al*. contained a recording corresponding to Or33b. Homotype DM3 coexpresses Or33b and Or47b, while homotype DM5 coexpresses Or33b and Or85a [29], which we incorporated manually by adjusting the odor response profile. When testing naturalistic inputs, we only chose odorants that elicited a strong response (≥40) to at least a single recorded homotype. This criterion resulted in odor response profiles for 39 homotypes to 96 odorants (+baseline firing rate). A keen reader might notice that neither of the datasets was comprehensive enough to cover all 58 homotypes available in the hemibrain dataset. For any missing measurements, we assumed no response and assigned zero values.

The functional group information was taken from Hallem *et al*. [25], while the odor valence information was acquired from the literature [6, 32–35].

## III. ALGORITHM OF COMPRESSED SENSING

Since its original inception by a group of mathematicians and engineers [36, 37], the compressed sensing has garnered substantial interest due to the apparent overcoming of the Shannon-Nyquist sampling theorem [38, 39], leading to its application in many different areas [40–42] For a compressible signal that can be mapped to a sparse representation, CS is employed to recover the original signal by solving for the sparsest solution of a set of under-determined linear equations. The first set of applications focuses on minimizing the quantity of measured input when the measurement process is expensive. Another set of applications recognizes the implementation of sparse representation for a highdimensional signal. The primary example would be various attempts at analyzing the signal processing inside a brain. The brain has been suggested to benefit from utilizing CS as a means to reduce the bandwidth and storage necessary for neural processes, including vision [43], olfaction [10, 17, 20–23], and others [44, 45], thus inferring high-dimensional signals from low-dimensional sensory input [46, 47].

Consider a set of linear equations

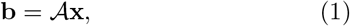

where **x** is an unknown *n*-dimensional vector, **b** is a known *m*-dimensional vector, and 𝒜 is a known *m* ×*n* matrix. If our goal is to find **x** that satisfies the equation for given **b** and 𝒜, the problem is trivial when the dimension of **b** is larger than the dimension of **x** (*m ≥ n*). However, when *m < n*, this is an under-determined (over-parameterized) problem, allowing infinitely many solutions for **x**. If our interest is to find 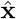, the sparsest solution of **x** subject to Eq. 1 such that the vector **x** has the smallest number of nonzero values, our problem becomes

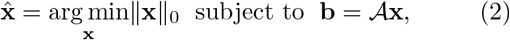

where ∥·∥_0_ is the cardinality or the *l*_0_ pseudo-norm that counts the nonzero elements in a vector such that ∥**x**∥_0_ = *K* is the sparsity of the vector **x**. However, minimizing the cardinality of **x** under a constraint corresponds to a non-convex optimization problem, which is computationally demanding. CS modifies Eq. 2 by considering an *l*_1_-norm minimization instead, which algorithmically simplifies Eq. 2 to a convex optimization problem:

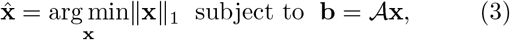

where 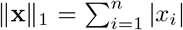. If the following two conditions are met, the solution of Eq. 3 is identical with that of Eq. 2, allowing one to recover the sparsest solution 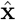 with a high probability [36, 39]. First, as the general rule of thumb, the dimension of **b** should satisfy a condition of *m > m*_*c*_ with *m*_*c*_ ≈ K log (*n/K*). Next, the sensing matrix 𝒜 must be incoherent, such that the columns in sensing matrix 𝒜 must be uncorrelated to each other, producing small inner products (see Appendix A).

We constructed the sensing matrix 𝒜 by taking a product of the normalized synaptic connectivity matrices 𝒞 ^PN−KC^ and 𝒞 ^KC−MBON^ (Fig. S1). That is, each column and row of 𝒞 ^PN−KC^ and 𝒞 ^KC−MBON^ is normalized, respectively. The *l*_1_-norm minimization was performed to generate the solution 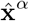 for a given MBON activity profile **b**^*meas,α*^ acquired from uPN expression to a specific odorant. We used Sequential Least SQuares Programming (SLSQP) optimizer implemented in the Scipy package constrained to **b** = 𝒜**x** with unconstrained variable boundaries.

When assessing the capacity of CS in odor identification for an odorant *α*, we evaluate the similarity between **b**^meas,*α*^ and 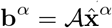, where 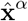 is the solution of Eq. 3 (see Appendix B).

## IV. THE CAPACITY OF COMPRESSED SENSING IN SIGNAL RECOVERY

We first demonstrate that CS can reliably recover the uPN activity profile from the MBON response profile through the sensing matrix defined by the product of two synaptic connectivity matrices (Fig. S1). Consider a situation depicted in Fig. 1A, where *Drosophila* whiffs the distinctive odor of an apple. The apple odor in the chemical space will be encoded into ORN/uPN activity profiles. Before an olfactory signal reaches the higher cortical area, the signal is integrated as it passes through higher-order olfactory neurons such as KCs and MBONs. Then, is it possible for the higher cortical area to infer a high-dimensional uPN activity from a low-dimensional MBON response profile and identify the odor code as an apple? It is not straightforward to gauge the perceptual differences between odors by simply comparing two MBON response profiles represented in 56-dimensional vectors (see the MBON profiles of methyl benzoate and ethyl benzoate in Fig S2, for instance, and the analysis given in Appendix C). Therefore, given the MBON response profile, we leverage CS to solve the inverse problem of finding the sparest representation of the odor in uPN activities instead. Ideally, the MBON activity expected from the solution of CS processed through the sensing matrix should reproduce what was given. Furthermore, provided that the odor code of an apple is clearly discernible from that of other odors, the odor representation of an apple at MBONs should be unique and discernible from others.

In CS, an MBON response profile and a uPN activity profile are translated to the vectors **b** and **x**, respectively. We build uPN activities (**x**^*meas,α*^) from the electrophysiological recordings [25, 26] for various odorants or odorant mixtures *α* and calculate the corresponding MBON response profile via the relation **b**^*meas,α*^ = 𝒜**x**^*meas,α*^. CS uses **b**^*meas,α*^ and 𝒜 to determine the sparest uPN representation for 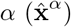. We next calculate the “residual” denoted by *r*_*α*_|_*β*_ (see Appendix B for the further detail), defined between the measurement (**b**^*meas,α*^) and the calculation 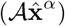 projected on itself 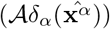 or other odor 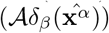 (Appendix B). The spectrum of residuals ({*r*_*α*|*β*_}), especially the gap between the self-residual (*r*_*α*|*α*_) and the cross-residuals (*r*_*α*|*β*_, *β* ≠ *α*), provides an insight into the differentiability of the odor *α* in the perceptual odor space (Fig. 2). If the self-residual is small and well separated from all other cross-residuals (*r*_*α*|*α*_ ≪*r*_*α*|*β*_ for all *β* ≠*α*), then the representation of odor *α* in the perceptual space is unique and easily discernible from others (Fig. 2, top). In contrast, if the self-residual is comparable to other cross-residuals (*r*_*α*|*α*_ ≃*r*_*α*|*β*_), it is difficult to differentiate the odor *α* from other odors, implying that the perceptual odor space of *α* is not well isolated (Fig. 2, bottom). Offering glimpses into the high-dimensional perceptual odor space, the spectrum of residuals {*r*_*α*|*β*_} allows us to assess the specificity of the odor *α* among other odors.

Embracing the criterion by Seki *et al*., who used either 30 spikes/s or 50 spikes/s as the detection threshold to demonstrate that a strong response to an odor is sparse [26], we define the strong response as the one evoking more than 40 spikes/s, analogous to the “winner takes all” (WTA) approach [19], and take into account only the odorants that elicit a strong response to at least one glomerular homotype in the experiments. The combined dataset contains the recordings from 39 glomeruli, which correspond to ∼80 uPNs (Fig. 3), in response to 96 different odorants encompassing diverse odor types that feature different functional groups and behavioral responses (see Table S1). Here, we assume that the activation of neurons is linearly correlated with equal gain in response to the odorants [48] and that uPNs extending from the same glomerulus receive identical input. Furthermore, we also assume that ∼40 uPNs out of 119 uPNs included in our circuit, which are not part of the measured 39 glomeruli, do not respond to the tested odorants.

**FIG. 3.**
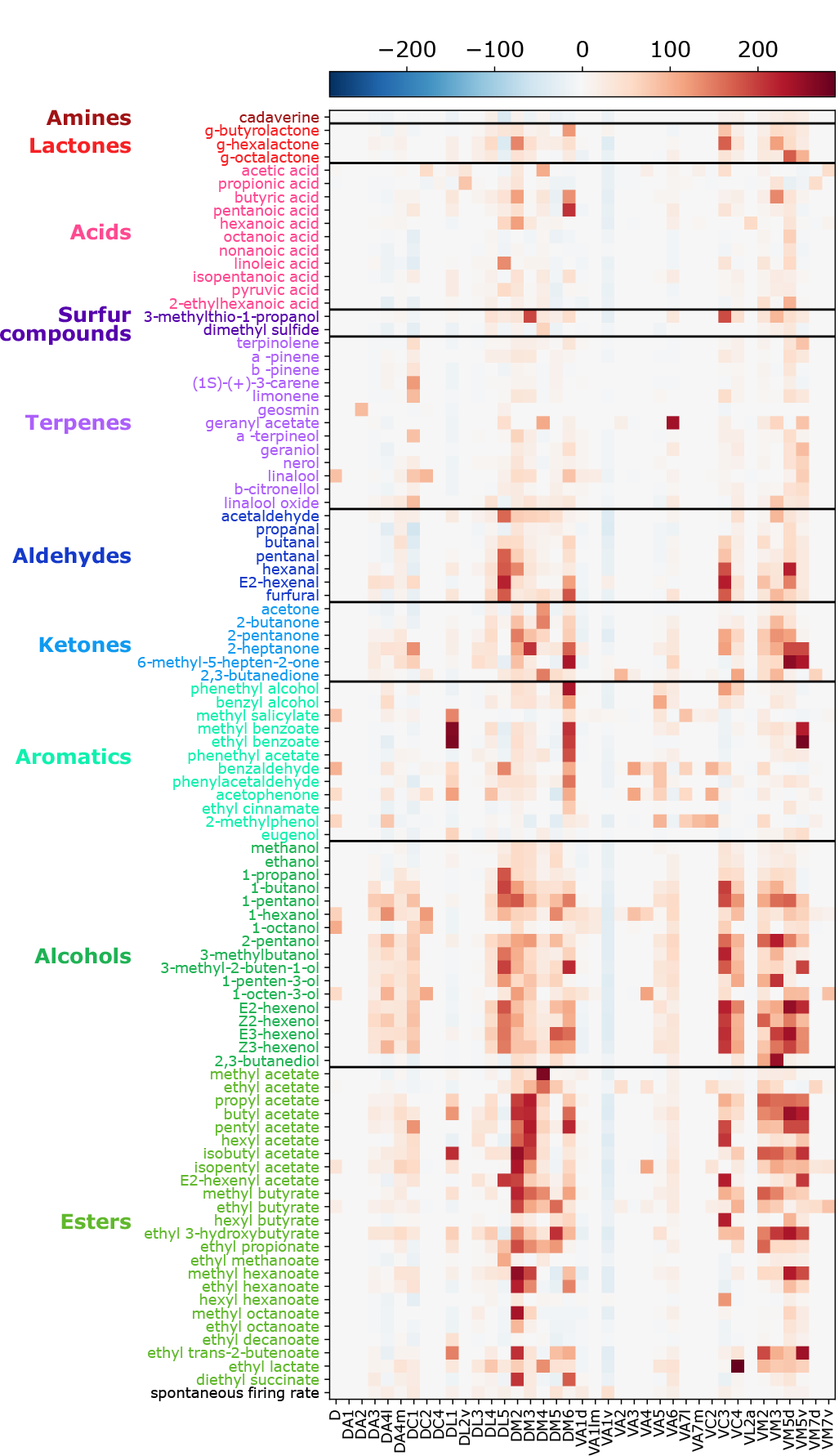
uPN activity profiles sorted by odorant functional groups. The electrophysiological recordings of 39 homotypes to 96 odorants (+ basal response labeled as ‘spontaneous firing rate’) where at least a single homotype exhibited a strong response (≥40 spikes/s). The labels are colorcoded based on the same odor categorization used by Hallem *et al*. [25] (Dark red: amines, red: lactones, pink: acids, purple: sulfur compounds, violet: terpenes, dark blue: aldehydes, blue: ketones, emerald: aromatics, green: alcohols, olive: esters)

Our CS framework correctly identifies 86 odorants out of 96 odorants (∼ 90%), indicating that 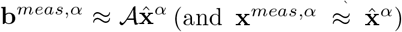, with *r*_*α*|*α*_ being minimal among all the residuals {(*r*_*α*|*β*_)} (Fig. 4. See Fig. S3 for an alternative demonstration of the result). The 10 odorants that failed our test are either alcohols or esters, which include 1-hexanol, 1-pentanol, 2-pentanol, cis-2-Hexen-1-ol (Z2-hexenol), trans-2-hexen-1-ol (E2hexenol), cis-3-Hexen-1-ol (Z3-hexenol), trans-3-hexen-1ol (E3-hexenol), propyl acetate, ethyl 3-hydroxybutyrate, and butyl acetate. For the statistical analysis of the residual spectrum for an odorant *α* ({*r*_*α*|*β*_}), we calculate the *Z*-score of the self-residual (*r*_*α*|*α*_) (see Appendix B). Many odorants exhibit significantly negative *Z*-scores, indicative of a large gap between *r*_*α*|*α*_ from *r*_*α*|*β*_ (*β*≠*α*) and a well-partitioned perceptual odor space for *α* (Fig. 5A).

**FIG. 4.**
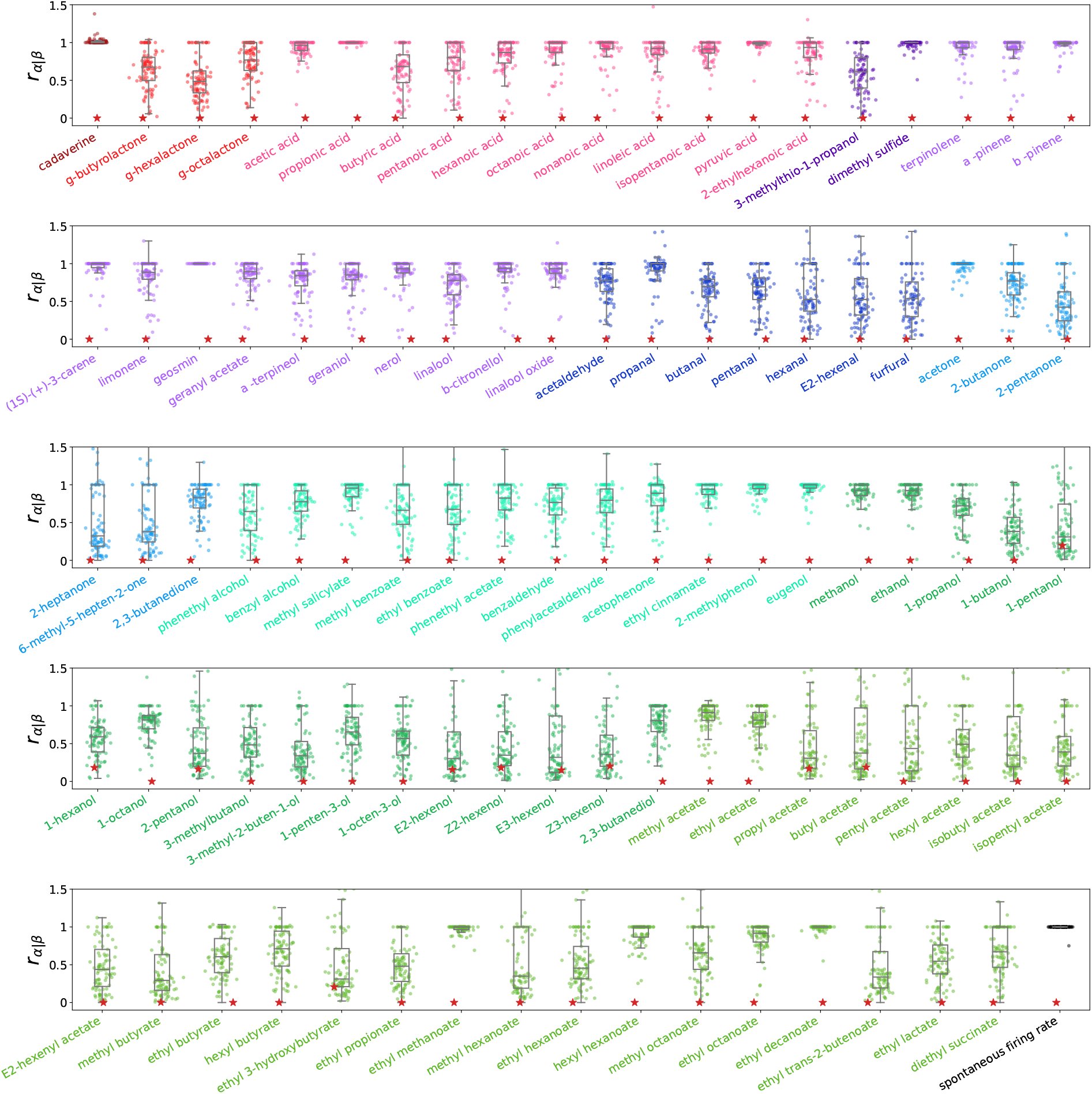
The residual spectra, {*r*_*α*|*β*_}, calculated for 96 odorants. The residual *r*_*α*|*β*_ calculated for an input odorant *α* against other odorants *β* denoted in the horizontal axis. Red stars denote the self-residual *r*_*α*|*α*_. The same color-codes as Fig. 3 are used.

**FIG. 5.**
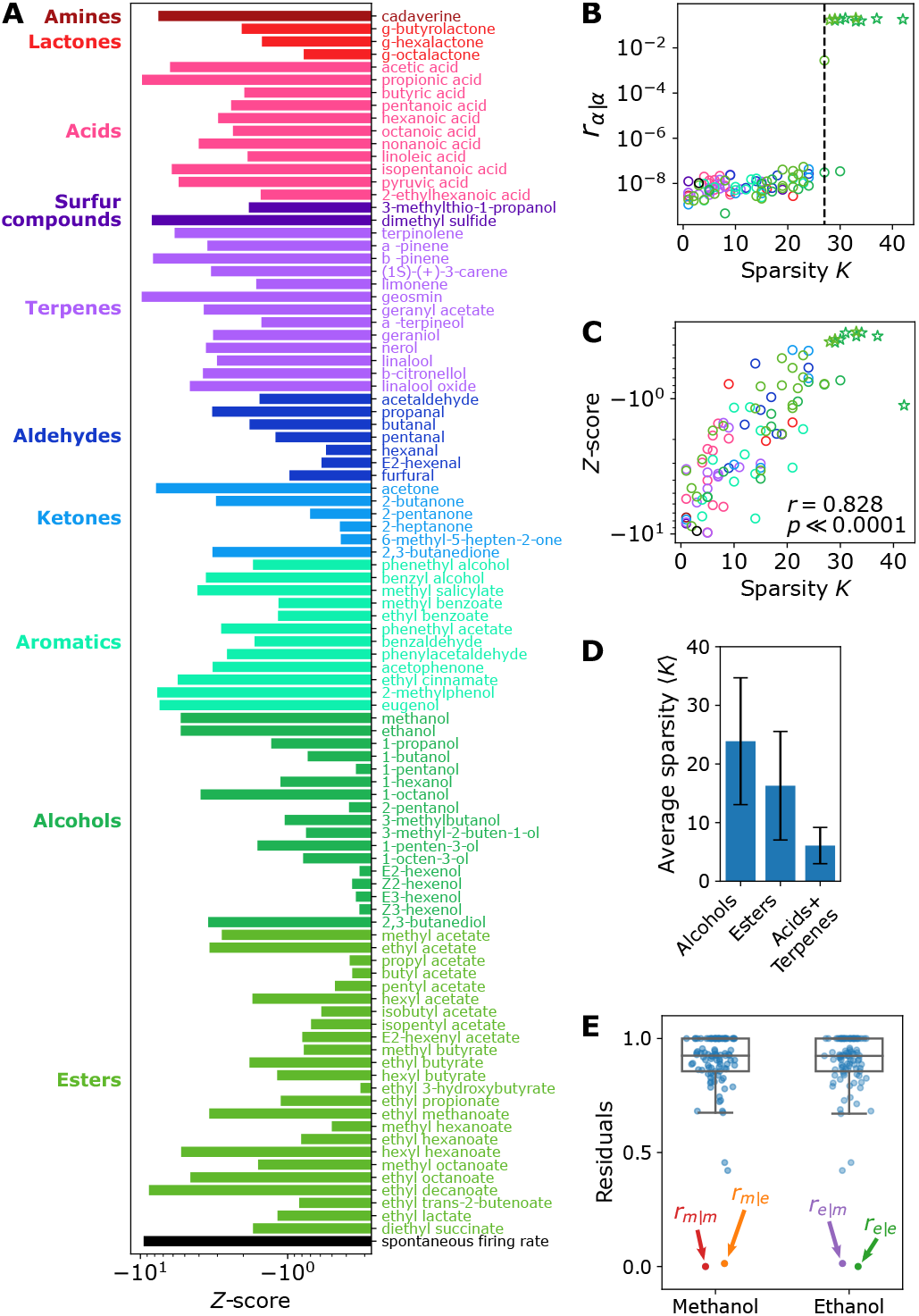
*Z*-score and sparsity *K*. (A) *Z*-scores of the self-residuals for 96 odorants calculated from Fig. S3. (B) Self-residual *r*_*α*|*α*_ versus the sparsity of **x**^*meas,α*^. The odorants whose self-residual (*r*_*α*|*α*_) is not minimal in the spectrum of residuals are marked with the star symbol. The black dashed line denotes the midpoint sparsity (*K* = 27) obtained by fitting the data to a logistic function. (C) *Z*-scores of *r*_*α*|*α*_ versus the sparsity (*K*) of **x**^*meas,α*^. (D) The average sparsity of the odor response profile for odorants that displayed strong responses (≥ 40 spikes/s) under the specified functional group. (E) Distributions of the residuals for *α* = MeOH and *α* = EtOH (*r*_*m*|*m*_ = *r*_MeOH|MeOH_, *r*_*m*|*e*_ = *r*_MeOH|EtOH_, *r*_*e*|*e*_ = *r*_EtOH|EtOH_, and *r*_*e*|*m*_ = *r*_EtOH|MeOH_).

Successful identification of odorant *α* hinges on the sparsity (*K*) of its input uPN activity profiles **x**^*meas,α*^ (Fig. 5B). An activity profile with a large *K*, which is found to correlate with the *Z*-score (Fig. 5C), gives rise to a large self-residual (*r*_*α*|*α*_) resulting in a *Z*-score with *Z* ≲ 0, which implies a non-optimal solution of CS. For accurate odor identification, the sparsity should be smaller than *K* = 27, a number close to the theoretical limit where our CS framework with the predetermined dimensions might experience the most difficulty in solving for the sparest solution (see Appendix A).

### A. Odor Identifiability Varies Across Odorants with Different Functional Groups

Odorants with certain functional groups, such as acids, terpenes, and aromatics, are characterized by *Z*-scores with *Z* ≪ 0 (Figs. S3 and 5), implying that the uPN activation profiles for these odorants are unique at least among the odorants we tested, and hence the corresponding perceptual odor space can be unambiguously discerned from others. In contrast, alcohols and esters elicit stronger responses over many uPNs [49], resulting in *Z* ≲ 0 (see Figs. 5A and S3). The average sparsity (⟨ *K*⟩) of the activity profiles for alcohols and esters is greater than that of acids and terpenes combined by 3−4 fold (Fig. 5D). The considerably high ⟨ *K*⟩ of alcohols and esters are likely linked to their poor odor identifiability by the *Drosophila* olfactory system. This finding is consistent with a previous study suggesting that ORs encoding alcohols and esters are broadly tuned, while another set of ORs that encode acids and amines are narrowly tuned [50]. The relatively poor odor identifiability of alcohols and esters, revealed from our CS framework, might also be related to the fact that the information necessary to identify alcohols and esters is processed through channels other than MBONs. Specifically, stereotyped spatial segregation of fruit-odor encoding PNs has been observed [11], which leads to stereotyped connectivity to LHNs [14, 51], suggestive of the existence of labeled lines [6, 12, 14, 26, 33, 35]. The odor identification and the subsequent decision-making process for alcohols and esters may heavily rely on the co-processing of information transmitted through both KCs and LHNs, as a sizable portion of LH projections converge with the down-stream projection from MB calyx [6, 27, 52].

Still, esters such as ethyl methanoate and ethyl decanoate are well discernible in the perceptual space. The same holds for methanol and ethanol, except when compared to each other. The odor identifiability assessed from our CS framework suggests that *Drosophila* may not be able to tell the difference between methanol and ethanol [25], as the corresponding values of self- and cross-residuals are almost identical to each other (*r*_*m*|*m*_ ≈ *r*_*m*|*e*_ and *r*_*e*|*e*_ ≈ *r*_*e*|*m*_. See Fig. 5E).

### B. The Perceptual Space of Odor Mixture and Olfactory White

In natural environments, most odors originate from complex mixtures of odorants. In humans, it is known that exposure to multiple odorants leads to the loss of the odor characterization of individual components [53–55], with a mixture consisting of an even greater number of odorants leading to a perceptual state called “olfactory white” [56]. This loss of odor differentiability is analogous to the loss in CS capacity, leading to a question of how simultaneous exposure to multiple odorants affects our CS framework. To test this hypothesis, we examine our framework against (i) random mixtures of a predetermined number of odorants and (ii) natural odor mixtures of fruit extract.

When multiple odorants are randomly mixed, how does the brain perceive the odor? At the glomerular level, it has been suggested that an odorant mixture elicits glomerular activities [57] that may or may not be identical to the sum of the activities of individual glomeruli [58–60]. Competitive binding of odorants to receptors is expected [60], and some odorants may be more dominant than others depending on the chemical and perceptual composition of the mixture [48]. Here we try to assess the odor identifiability from the vector **x**^*meas,α*^ encoding a natural odor whose precise chemical composition is unknown. First, we estimate the sparsity of the uPN activity profile associated with natural odors since the challenges encountered by the olfactory system translate to the difficulty of signal recovery via CS. We compare the results from single odorants and fruit extracts, the latter of which are available from Hallem *et al*. in the form of electrophysiological recordings on 23 glomeruli [25]. To make a fair comparison of fruit extracts against the mixture of single odorants, we analyze the responses to the single odorants using only 23 glomeruli, which are extended to the 47 uPNs, as in the actual measurement (see Fig. 6A). The chemical composition of fruit odors is expected to be diverse, and we observe a large variation in the sparsity among fruits. The sparsities of some fruits (e.g., banana) are high, to the point that their odor identifiability through CS is anticipated to be low. There is a chance that the sparsity reported here is underestimated since other sets of glomeruli not recorded in the experiment may have activated. Furthermore, a natural odor, whose chemical composition is complex in reality, could display a small *K* when it is encoded by a small set of broadly-tuned ORs [61].

**FIG. 6.**
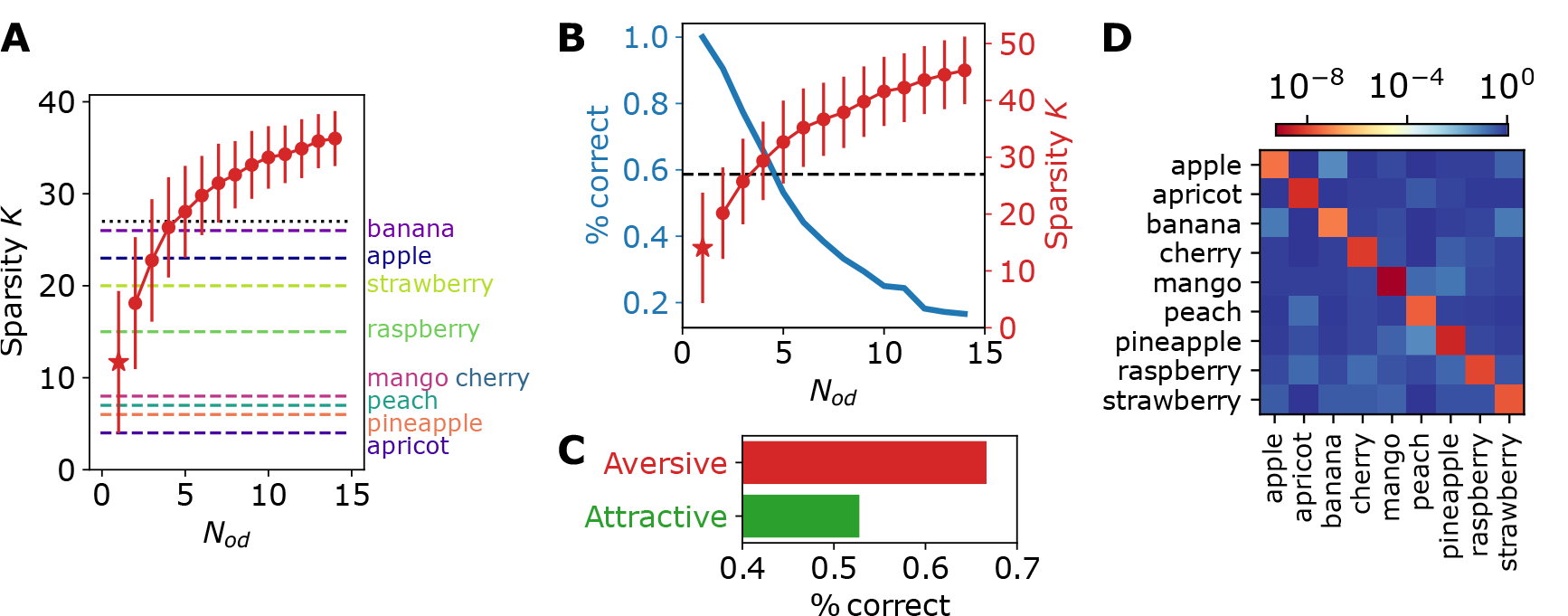
Analysis of odor mixtures. (A) The average sparsity of randomly sampled odor mixture with different sample size *N*_*od*_ (red dots) and the sparsity of **x**^*meas,α*^ when *α* = natural mixtures (dashed line). The sparsities are counted based on the 23 glomeruli whose responses were recorded in both the single odorant experiment and the fruit odor experiment for consistency. The red star denotes the average sparsity for single (*N*_*od*_ = 1) odorant exposure. The error bars indicate the standard deviation. The dotted black line corresponds to *K* = 27 in Fig. 5B. (B) The input signal recovery (blue line) and the corresponding average sparsity (red dots) of a random mixture of odorants with different sample size *N*_*od*_. The output is from the combined dataset measuring 39 glomeruli against 86 odorants whose self-residual (*r*_*α*|*α*_) was minimal. The red star denotes the average sparsity for single (*N*_*od*_ = 1) odorant exposure. The red errorbar indicates the standard deviation. The dashed line corresponds to the midpoint sparsity (*K* = 27). (C) The probability of successful input signal recovery for the mixture consisting of two attractive odors (green) or two aversive odors (red). (D) The matrix of the residuals, *r*_*α*|*β*_, calculated based on fruit odors. Each row depicts the spectrum of the residuals of an odor input *α* with respect to other fruit odors *β* (*β* = apple, apricot, …) denoted in the horizontal axis.

For random odorant mixtures, we directly compared vectors 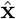 and **x**^*meas*^ to assess the accuracy of the signal recovery instead of calculating the residual spectra. Specifically, we assumed that the recovery was successful if the cosine distance between the two vectors, defined by 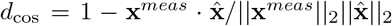, is smaller than 0.05. From the 86 odorants that the CS framework accurately identified, we randomly sampled the odorants to create 500 random mixtures with size *N*_*od*_, linearly combined the uPN activity profiles of individual odorants in each mixture, and performed CS to infer 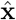. As expected, we observe a steady decline in the chance of successful recovery of the original signal **x**^*meas*^ as the number of odorants composing a mixture (*N*_*od*_) increases, with the chance dropping below 20 % for *N*_*od*_ *>* 10 (or *K >* 40) (Fig. 6B). When many odorants are mixed at comparable concentrations, the signal recovery becomes less effective under our CS framework.

However, the nature of the random odorant mixture can vary when the mixture is restricted to a specific valence. We select nine attractive odorants and seven aversive odorants, all of which are well characterized in terms of *Drosophila* behavioral response [6, 14, 32–35, 62, 63] (see Table S1) and create combinations of attractive or aversive odors by sampling two odorants from each category. We find that the capacity for signal recovery is significantly poor for the mixture of attractive odors compared to the mixture of aversive odors (Fig. 6C). This may be associated with the generally poor odor identifiability of alcohols and esters (Fig. 5A) that constitute many attractive odorants. While our result may not be directly translatable to the odor perception of *Drosophila in vivo*, the functional consequence of loss in the identifiability of an odor mixture seems analogous to the notion of “olfactory white.”

In the case of natural mixtures (fruit extracts), we associated an activity profile for a fruit odor with a percept, such that *α* designates a type of fruit (e.g., apple). The measurements were made in response to the concentrated extracts from nine fruits (apple, apricot, banana, cherry, mango, peach, pineapple, raspberry, and strawberry) diluted to 10^−2^ volume per volume (v/v) with H_2_O. Based on the results from random odorant mixtures, the recovery of fruits such as apples and bananas whose uPN activity profile has high sparsity does not seem guaranteed (Fig. 6A). However, the spectrum of residuals is still specific enough to resolve the corresponding odors among a list of fruits, suggesting that their perceptual odor spaces in the brain of *Drosophila* are still well separated from other fruits (see Figs. 6D and S4).

### C. Odor Perception May Depend on the Odorant Concentration

So far, our analysis has been conducted using electrophysiological recordings of odorants at a relatively high concentration. Both the recordings provided by Hallem and Carlson [25] and Seki *et al*. [26] employed 10^−2^ dilution with either H_2_O, paraffin oil, or mineral oil (except for geosmin, which used 10^−3^ dilution) [26]. It is, however, known that the activity profile of ORNs and PNs are concentration-dependent [18, 25], and so is the observed hedonic valence of *Drosophila* to a given odorant [62, 64]. Similar concentration-dependent behavioral responses have been observed in *C. elegans* [65].

Fig. 7A summarizes the electrophysiological responses of glomeruli to a small subset of odorants at varying dilution levels (10^−8^, 10^−6^, 10^−4^, 10^−2^) [25]. Some odorants evoke strong responses at low dilutions while others do not, consistent with the studies on odorant-dependent odor detection thresholds [66–68]. Odorants at unusually low concentrations may still be perceivable through a combination of specialized olfactory processing [69, 70]; however, for consistency, we limit our study to dilution levels where there is an activation from at least a single neuron (see Fig. 7A and S5A). All eight odorants failed to evoke strong responses at 10^−8^ dilutions.

**FIG. 7.**
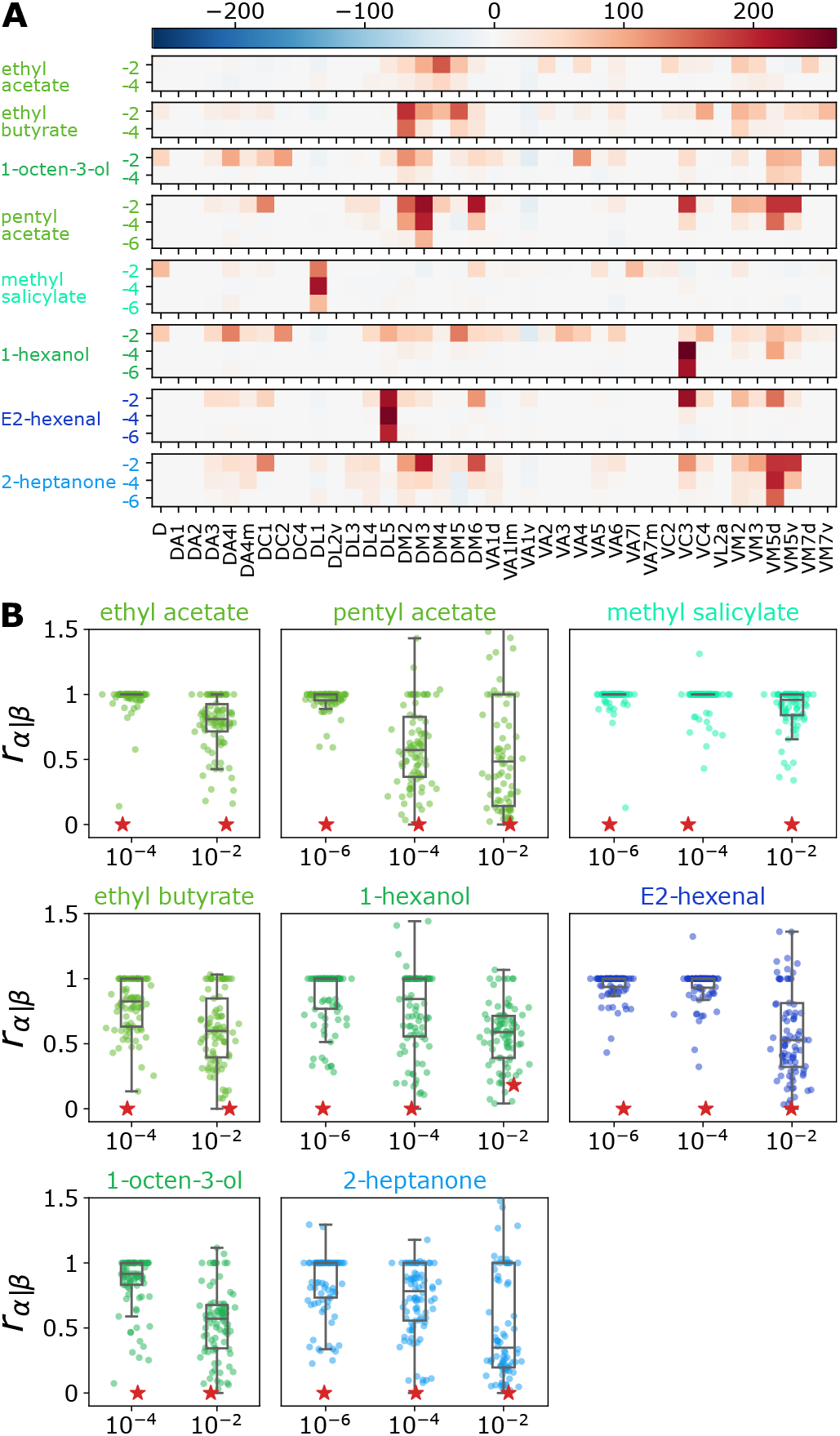
Concentration-dependent perception of odor. (A) Electrophysiological recordings of uPN activity profiles in response to select odorants measured at different dilution levels (−2, −4, and −6 on the vertical axes signify the dilutions of 10^−2^, 10^−4^, and 10^−6^, respectively). Only those that elicited a strong response to at least one homotype are plotted. (B) The distributions of *r*_*α*|*β*_ for odorants at different dilution levels. Red stars denote the self-residual *r*_*α*|*α*_.

When performing CS, we included all the previously tested odorants, which were measured at 10^−2^ dilution while replacing the odorant *α* with different dilution levels. 1-hexanol, an odorant that evaded our CS framework at a high concentration, became identifiable at lower concentrations, presenting a minimal self-residual (see Fig. 7B). A significant reduction in the sparsity *K* at a lower odorant concentration (see Fig. S5A) helps CS accurately recover the uPN activity profile. Our finding also translates to natural mixtures. The high identifiability of fruit odor at 10^−2^ dilution drops significantly for undiluted fruit extracts (see Fig. S5B).

Significant changes are observed for the residual spectra of the same odorant at different concentrations (see Fig. 7B). For example, the residual distributions of pentyl acetate and trans-2-hexenal exhibit a drastic change from 10^−6^ to 10^−4^ and from 10^−4^ and 10^−2^ dilutions, respectively. Physiologically, a drastic shift in the residual distributions implies considerable changes in uPN activity and the corresponding MBON response profiles (see Fig. S5C). We surmise that this signifies an alteration of the odor percept in the perceptual space to the point that the odorant under two concentrations may be perceived as different odors. For ethyl butyrate, our calculation shows a concentration-dependent shift in the residual distribution, explaining the conflicting information on the *Drosophila* behavioral responses reported in the literature [6, 62, 71]. These results are reminiscent of the changes in the qualitative properties reported in human odor characterization experiments.

Taken together, the sparse encoding of odor to the receptor space is a prerequisite for odor identifiability under our CS framework, and an odor under two vastly different concentrations may occupy disparate perceptual spaces, leading to a different odor percept.

## V. DISCUSSION

The neurons and the synaptic connectivity associated with the *Drosophila* olfactory system are akin to the nodes and weight matrices of artificial neural networks (ANN). Whereas an ANN dynamically trains the weight matrix to gain functionality, our system operates on a pre-configured synaptic weight matrix. The synaptic characteristics observed from the *Drosophila* olfactory system, such as the seemingly randomized synaptic interface between uPNs and KCs (Fig. S1) and the sparsity of uPN activity profiles (Fig. 3), are reminiscent of the prerequisite for CS. Hence, we have applied CS to the physiological circuits of the *Drosophila* olfactory system to explore its capability of inferring the identity of the sensed odorant from its representation in the inner brain. By leveraging the relations of neural codes in the receptor space to the perceptual odor space represented by the MBON response profiles, we study the differentiability of an odor stimulus in the perceptual odor space and provide a conceptual explanation of perceptual phenomena such as olfactory white and concentration-dependent odor perception.

As previous studies have acknowledged, whether or not CS is implemented in the *Drosophila* brain is unknown [17, 72]. There is no direct evidence that the higher cortical area effectively leverages the randomization of the PN-KC interface and that there exists a necessary circuitry to perform *l*_1_-norm minimization, although there are suggestions for the implementation of sparse coding in real organisms [46, 73]. Nonetheless, it is fascinating to discover that the *Drosophila* olfactory system exhibits the structural prerequisites for CS. We hypothesize that for the downstream neural circuitry beyond the odorant-OR interface, the compression of the information for chemical stimuli through the effectively randomized sensing matrix is beneficial for conserving signal integrity while reducing the bandwidth and resources necessary for signal transmission to the higher cortical area. It is also interesting to note that a randomly pre-wired network has shown to be more cost-effective with improved accuracy and network trainability in the recent computational development in ANN [74].

While not explicitly considered in this study, other neuronal architectures of the olfactory system are known to play critical roles in signal processing. The most notable are the peripheral interactions [60] present in the second and third-order olfactory neurons. By suppressing the sensitivity of PNs to ORN inputs, lateral inhibitions [75] can allow the encoding of a broader range of concentrations [61], sparsifying the olfactory signal. Mixing highly overlapping odorants that activate similar PNs is known to promote lateral inhibition [76], which could enhance the chance of odor identification even if the uPN activity profile is over-saturated in our framework. Additionally, noise filtration is implemented through inhibitory circuits composed of various antennal lobe local neurons (LNs) [77–79], GABAergic inhibitory PNs (iPNs) [13, 51], anterior paired lateral neurons (APLs) [6, 80, 81], etc. The regulations of olfactory signals across glomeruli by LNs or KCs by APLs lead to noise reduction and sparsification of odor representation [24, 75], both of which should improve the reliability of our CS framework. While we find that the 86 odorants maintain a respectable level of recovery even in the presence of a significant amount of noise (see Fig. 11A), the reduced noise can further sparsify the vector **x**^*meas*^ and ease recovery and identification of the correct activity profile in theory, especially for an odor mixture where a low sparsity of **x**^*meas*^ is not guaranteed. Moreover, ORN-PN sensitivities are glomerulus-specific [82], indicating differential gain control. The ORN response to an odor is nonlinear [18, 25], and a more sophisticated and physiologically correct model for the firing rate could have been used [17]. Nonlinearity prevalent in the olfactory system [18, 61, 77] plays a significant role in olfactory processing [17, 22, 80], and many of these regulatory circuits display odor-type specificity [13, 61].

The temporal dynamics of the signal are critical for olfactory processing as well. For odor perception, timevarying changes in odor concentration play a pivotal role in odor identification, decision-making, and odor-guided navigation [16, 83, 84]. Other interesting psychophysical phenomena such as olfactory fatigue (peripheral olfactory adaptation) [85], require temporal information to process. Additionally, the temporal dynamics are closely related to the primacy coding hypothesis, which predicts earlier olfactory signals to govern the overall odor identity [16], thereby significantly reducing the difficulty of signal recovery at higher concentrations under our CS framework if implemented. Whereas we did not implement primacy coding due to the limitation in data, it may still be possible to infer the temporal characteristics from the concentration-dependent changes in the uPN activity profile as they highlight ORs most sensitive to a given odorant which should correlate with the order of activity. While we believe that the primacy coding is an effective principle behind concentration-invariant odor identification, we suspect that the less sensitive and therefore delayed ORN responses may also contribute to the odor perception as characterized by the concentration dependent quality changes observed in humans.

Lastly, our framework does not consider the anatomy and physiology behind synapses and neurons, which have been shown to contribute to complex olfactory computation such as independent gain control [86]. The locations of presynaptic terminals can be crucial in multimodal sensory processing, and the differentiation between axodendritic and axo-axonic connections is necessary for spatiotemporal dynamics [87].

In this work, we explored the concentration-dependent changes to projected perceptual space by entirely attributing it to the changes in the odor codes (**x**^*meas*^). Regarding the concentration-dependent valence, the aversion at high concentration is mediated by the signal through LH that defines the innate responses, and the innate responses are in turn modulated by the output from MB calyx [26]. In light of classical conditioning experiments on *Drosophila* [88, 89], which demonstrated the dramatic change in the *Drosophila* behavior in response to electric shock, it can be surmised that behavioral responses generally depend on both the sensory processes of odor identification and the learned responses. We assume that modification in behavioral response (or decisionmaking) may result from the multimodal integration of various sensory inputs not covered in this study.

Taken together, we demonstrate the capacity of the CS framework to recover high-dimensional uPN activity profiles from low-dimensional *barcodes* at MBONs in the *Drosophila* olfactory system. In fact, partial information on the MBON response profile still suffices to recover the corresponding uPN activity profile for many odorants (Appendix D), suggestive of much room for more efficient data compression and retrieval. It will still be of great interest to explore the effects of incorporating the various neuronal features listed above into the basic structure of our CS framework.

## ACKNOWLEDGMENTS

All original data and codes are deposited and publicly available at https://github.com/kirichoi/DrosophilaPerception. This study was supported by KIAS Individual Grants CG077002 (K.C.), CG076002 (W.K.K.), and CG035003 (C.H.). We thank the Center for Advanced Computation in KIAS for providing the computing resources.

## Appendix A: Conditions for solving the inverse problem through compressed sensing

We performed a detailed analysis of the synaptic connectivity between the layers of olfactory neurons to check whether it conforms to the criteria for successful recovery of the sparsest solution 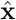 through CS. Unlike the previous applications of CS to the olfactory system [17, 22, 23], where the true dimension of the odorant space is unknown, the dimensions of input and output vectors are well defined in our problem.

First, CS requires that the size of the input (*m*, the dimension of **b**) be reasonably large compared to the dimension (*n*) and the sparsity (*K*) of the target vector to be recovered (**x**). According to the synaptic connectivity provided by *Drosophila* hemibrain connectome [5], out of 162 uPNs, only 119 uPNs innervate MB calyx and connect to KCs [14]. These *n* = 119 uPNs that synapse with at least one of ∼2000 KCs can be traced down to *m* = 56 MBONs based on the synaptic connectivity. We find that *m > m*_*c*_ ≃ *K* log (*n/K*), the empirical criterion of CS for a successful recovery of the 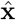 [90, 91], is satisfied for all values of *K* (Fig. 8A).

**FIG. 8.**
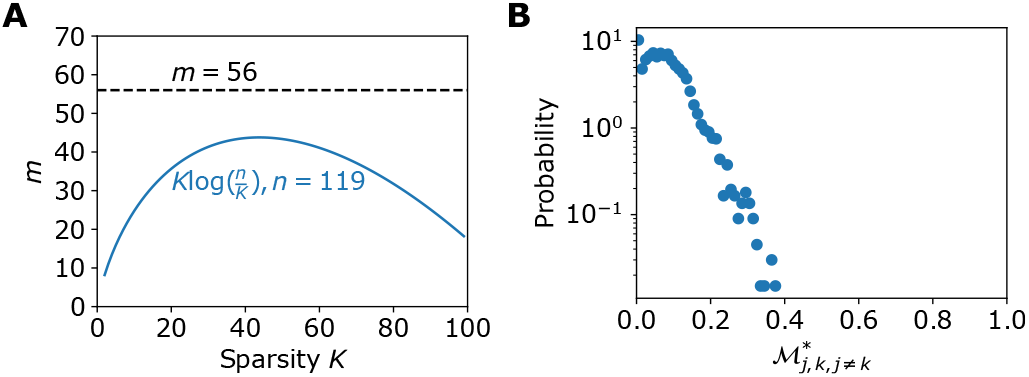
Criteria required for successful signal recovery using the CS framework. (A) A plot illustrating the gap between the dimension of MBON (*m* = 56) and the *K*-dependent curve *K* log (*n/K*) for *n* = 119. (B) The distribution of the pairwise correlation 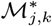 between the connectivity matrices 𝒞^PN−KC^ and 𝒞^KC−MBON^.

We also approximated the incoherence criterion of the sensing matrix 𝒜 constructed from two connectivity matrices, 𝒞^PN−KC^ and 𝒞^KC−MBON^, between uPNs, KCs, and MBONS. In the original framework of CS, the in-coherence of the sensing matrix 𝒜 is defined through a value [92, 93]

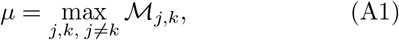

where

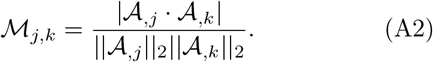

Here, 𝒜_,*i*_ denotes the *i*-th column vector of 𝒜. The value *μ* ranges from 0 to 1, with 0 indicating complete incoherence. Then, reconstructing a *K*-sparse vector 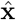 becomes highly probable under the following condition [93]:

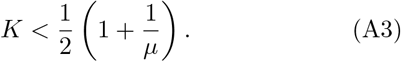

From the sensing matrix 𝒜= (𝒞^PN−KC^ · 𝒞^KC−MBON^)^⊺^ and Equation A1, we obtain *μ ≈*1, which is the theoretical maximum. Based on this result, the reconstruction of the vector 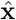 through *l*_1_-norm minimization seems infeasible. However, the requirement for incoherence defined in Eq. A1 is often considered too stringent to achieve, given that *μ* = 1 is obtained even when a single pair of columns in the sensing matrix 𝒜 are identical. As a result, several guidelines to relax the original criterion for incoherence have been suggested [92, 94], requiring that the sensing matrix be only *effectively incoherent*.

In practice, it has been suggested that the recovery of a *K*-sparse vector is achievable with a high probability from asymptotically small *μ* as the size 𝒜 of increases [93, 94]. Brunton and Kutz have suggested a more relaxed condition [39]. When the sensing matrix 𝒜 is a product of a measurement matrix 𝒮and a sparsifying basis Ψ, i.e., 𝒜= 𝒮Ψ, a small scalar product between the rows of and the columns of Ψ is a good indication that *l*_1_-norm minimization will recover 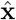, which replaces the matrix ℳ_*j,k*_ (Eq. A2) with

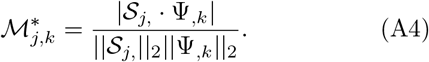

While our 𝒞 ^PN−KC^ and 𝒞 ^KC−MBON^ are not analogous to the measurement matrix and the sparsifying basis by definition, the suggested requirement is mathematically equivalent within the scope of successful *l*_1_-norm minimization subject to **b** = 𝒜**x**. When we adopt 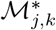 to approximate the incoherence criterion, a large portion of 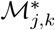 are zero or close to zero (Fig. 8B), and about 96% of the values are below 0.2, a theoretical limit which would allow CS to recover a target vector **x** with the sparsity of *K* = 3 (Eq. A3), which corresponds to an approximate average uPNs per glomerulus.

Restricted isometry property (RIP) is often cited as the rule of thumb when it comes to solving the inverse problem, but checking for RIP is often prohibitively expensive [93] and unnecessary in practical applications [94]. It has, indeed, been demonstrated that CS is possible even under weaker conditions [94, 96]. We find that the sensing matrix of the *Drosophila* olfactory system is effective in recovering 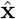 for all practical purposes.

## Appendix B: Assessing the odor identifiability by means of residual spectrum

The specificity (identifiability or differentiability) of an odor *α* against other odors is assessed by evaluating the similarity between (i) the *measurement* and (ii) the *calculation* of the MBON response profile. (i) The measured MBON response profile, denoted by **b**^*meas,α*^, is acquired using the electrophysiological recording of uPN activity profile **x**^*meas,α*^ and the sensing matrix 𝒜 via the relation **b**^*meas,α*^ = 𝒜**x**^*meas,α*^. (ii) A MBON response profile of the odor *α* can be calculated from 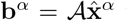 using the solution of the CS algorithm, i.e., 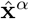 from Eq.3.

To assess the differentiability of the odor *α* against other odors {*β*}, we adapted the sparse representation for classification (SRC) [39, 95] and considered a function 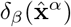 that filters the uPN activity profile calculated for the odor *α* against the uPN activity profile measured for the odor *β*. Specifically, the filtering function 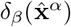 for the *i*-th component of the uPN activity profile is defined as

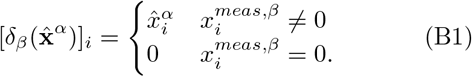

Note that the self-filtering yields the original profile, i.e., 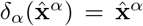. Along with Eq.B1, we defined *r*_*α*_ |_*β*_, the (relative) residual between the measurement (**b**^*meas,α*^) and the calculation 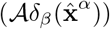 over all test odors *β* including *β* = *α*, which enables us to assess the specificity of the odor *α* against all other odors {*β*}:

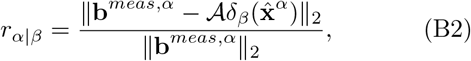

where 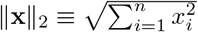.

Ideally, 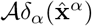 must be identical to **b**^*meas,α*^, i.e., *r*_*α*|*α*_ = 0, but in practice, we find *r*_*α*|*α*_ ≳ 0. However, if *r*_*α*|*α*_ ≪ *r*_*α*|*β*_ for all *β ≠ α*, then the odor represented by *α* can be considered specific and easily discernible from other odors. In contrast, when there are many odors, say *β*_1_, *β*_2_, …, satisfying 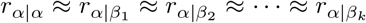, it means that the odor space (perceptual space) of *α* is not well isolated from that of *β*, suggesting ambiguous sensing of odor *α*. In this case, discerning the odor *α* from other odors *β*_1_, *β*_2_, …, *β*_*k*_ is challenging for *Drosophila*.

To assess the spectrum of the residuals for the odor *α*, we calculated the *Z*-score of *r*_*α*|*α*_ defined as

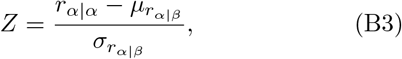

where 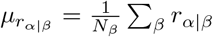 and 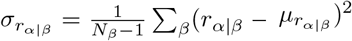. Therefore, a *Z*-score with *Z* ≪ 0 implies that the value of the self-residual (*r*_*α*|*α*_) is statistically well separated from the cross-residuals (*r*_*α*|*β*_), and the odor *α* is easily discernible from other odors.

## Appendix C: Analysis of MBON response profiles using Euclidean distances

In this study, we utilize CS to discuss the differentiability between the MBON response profiles of various odorants. In principle, one could perform a similar analysis by calculating the distances between MBON response profiles. We, however, find that the results from such an analysis on high-dimensional vectors are often limited by the issues associated with the curse of dimensionality [46]. We find that the recovery of the uPN activity through the CS algorithm is better suited for odor differentiability than other standardized methods.

Consider a pairwise distance matrix between MBON response profiles. Fig. 10A shows the distribution of the scaled Euclidean distances between uPN activity profiles, 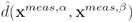, where

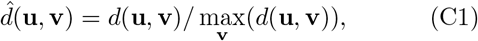

with 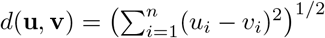 and 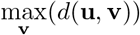 being the maximum among the distances from the vector **u** to other vectors **v**. The clear gap between the self-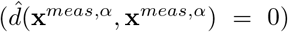 and cross-distance 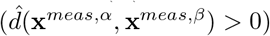 found in the uPN activity profiles is no longer observable in the MBON response profiles 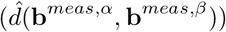. This suggests that the Euclidean distance and its derivative measures are inadequate to characterize the highly integrated signal at MBONs (Fig. 10A, bottom). We observed a similar trend when we used other metrics such as cosine distance.

**FIG. 9.**
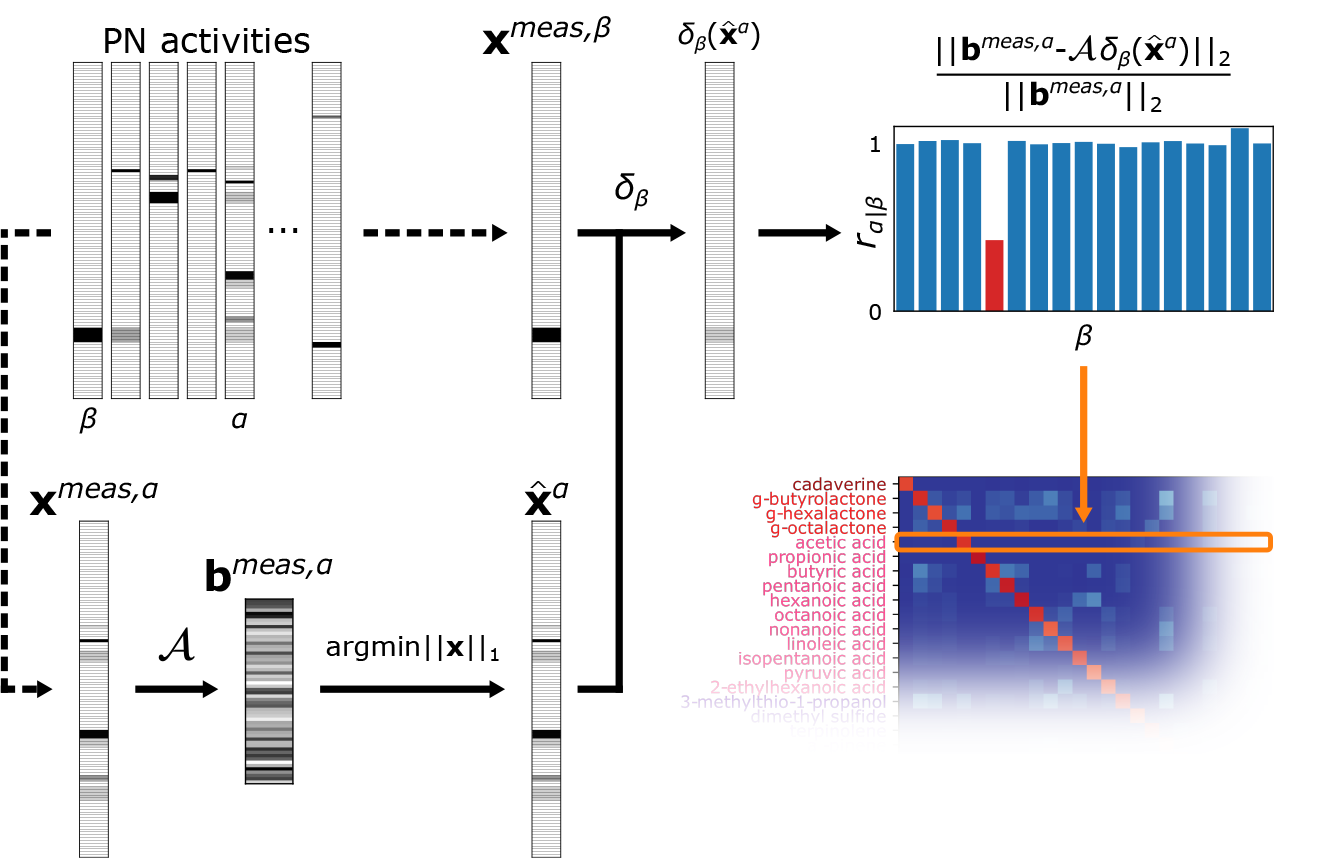
Calculation of residual spectrum. uPN activity profile is used to calculate the input MBON response profile **b**^*meas,α*^, where *α* is an odorant. *l*_1_-norm minimization is performed to determine the solution 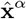 from a set of underdetermined linear equations, which is compared with the input uPN activity profile 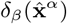 filtered based on the expected uPN activity from another odorant *β*, **x**^*meas,β*^. Adapting the framework of sparse representation for classification (SRC) [39, 95], we calculate the residuals *r*_*α*|*β*_ for a given input odorant *α* and reference odorant *β*. The residuals for an odorant *α* against all other reference odorants {*β*}, *r*_*α*|*β*_, are calculated.

**FIG. 10.**
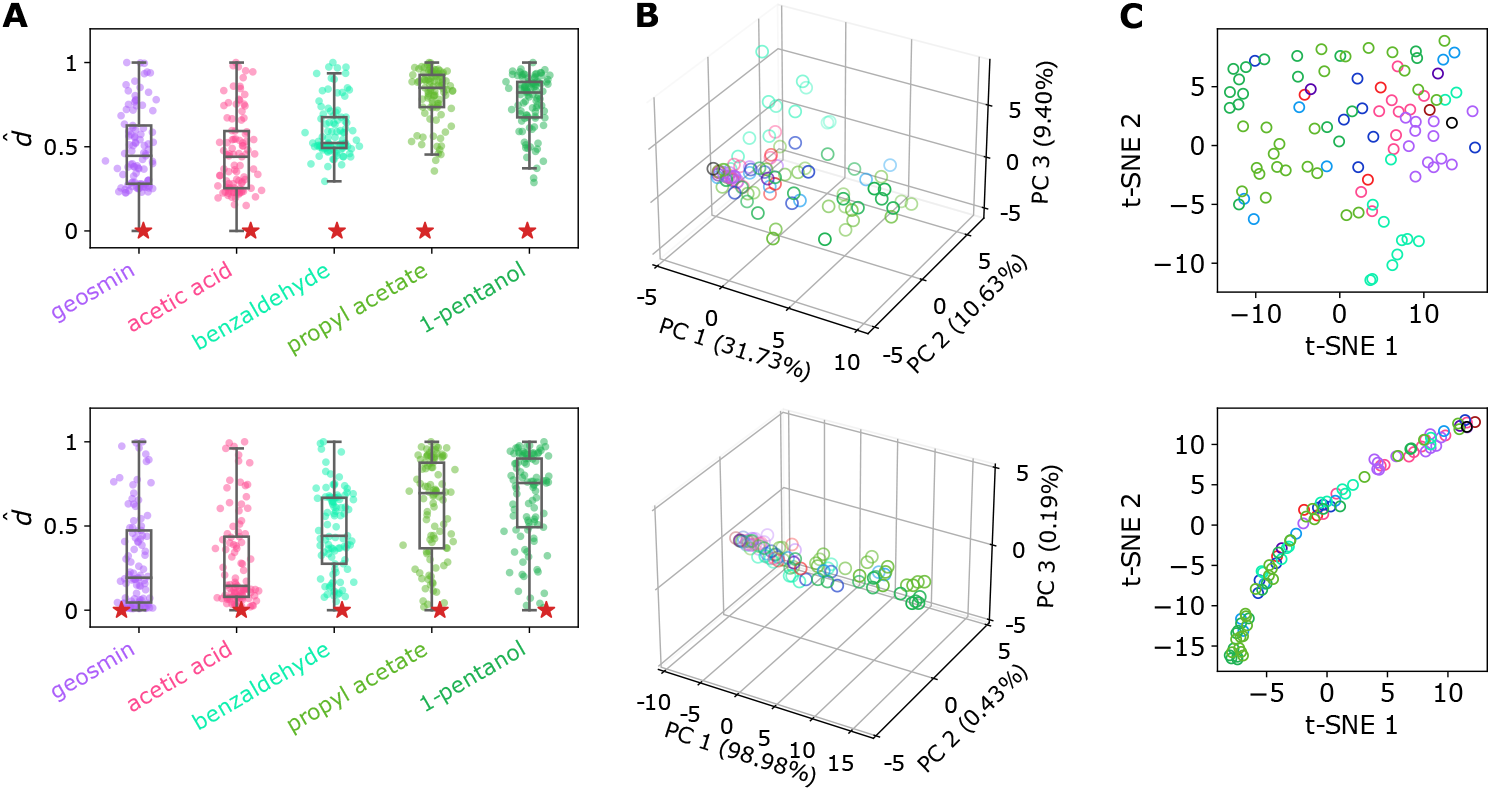
Analysis of MBON response profiles using the Euclidean distances. (A) The distribution of scaled Euclidean distances between (top) uPN activity profiles and (bottom) MBON response profiles in response to select odorants. (B) The output of PCA projecting (top) uPN activity profiles and (bottom) MBON response profiles to 3-dimensional space, along with the proportion of variance explained by each principal component. (C) The output of t-SNE projecting (top) uPN activity profiles and (bottom) MBON response profiles on two dimensions.

Incidentally, separation of the odor space by physio-chemical features of odorants and OR/PN activities have previously been shown using principal component analysis (PCA) and t-distributed stochastic neighbor embedding (t-SNE) [18, 25, 62, 63]. We performed PCA and t-SNE on a standardized matrix of MBON response profiles, in which each column of the matrix that represents MBON response profiles has zero mean and unit variance. PCA was performed using singular value decomposition (SVD), and t-SNE was performed using a perplexity of 15 after initializing with PCA. The result of PCA indicates that, unlike uPN activity profiles, many odorants are hard to differentiate using MBON response profiles (Fig. 10B), where the proportion of variance explained by each principal component quickly diminishes after the first principal component (Fig. 10B, bottom), suggestive of no clear separation in the odor representation. The lack of a clear separation in the component space is also found in the analysis using t-SNE (Fig. 10C).

## Appendix D: Odor identification using partial information from MBONs

Our CS framework reconstructs the uPN activity from the response profile of 56 MBONs, where we assume that the MBON response profile reflects the information transmitted to the higher cortical area. Here, we consider a hypothetical situation where only partial information from the MBONs is available to the higher cortical area, perhaps due to some stereotyped connectivity, intrinsic stochasticity, and limited access to MBONs at a given timeframe in animal perception, and ask whether the information encoded by a subset of MBONs is still enough to accurately infer the uPN activity profile [10]. For each of the 86 odorants our CS framework correctly identified, we randomly sample the MBON response profile 10 times with a sampling size *N*_MBON_ and examine whether at least one of the samples led to a correct identification. We find that a large number of odors can still be identified by using only a handful of MBONs, with the success rate improving with the sampling size *N*_MBON_ (see Fig. 11B). We observe that the compressibility of information necessary to identify the odor varies with the functional group of the odorants. In particular, alcohols and esters are the first to fail as the size of *N*_MBON_ decreases (see Fig. 11C). Odors from other functional groups (e.g., acids and terpenes) retain their accuracy even if the size of *N*_MBON_ is significantly small.

**FIG. 11.**
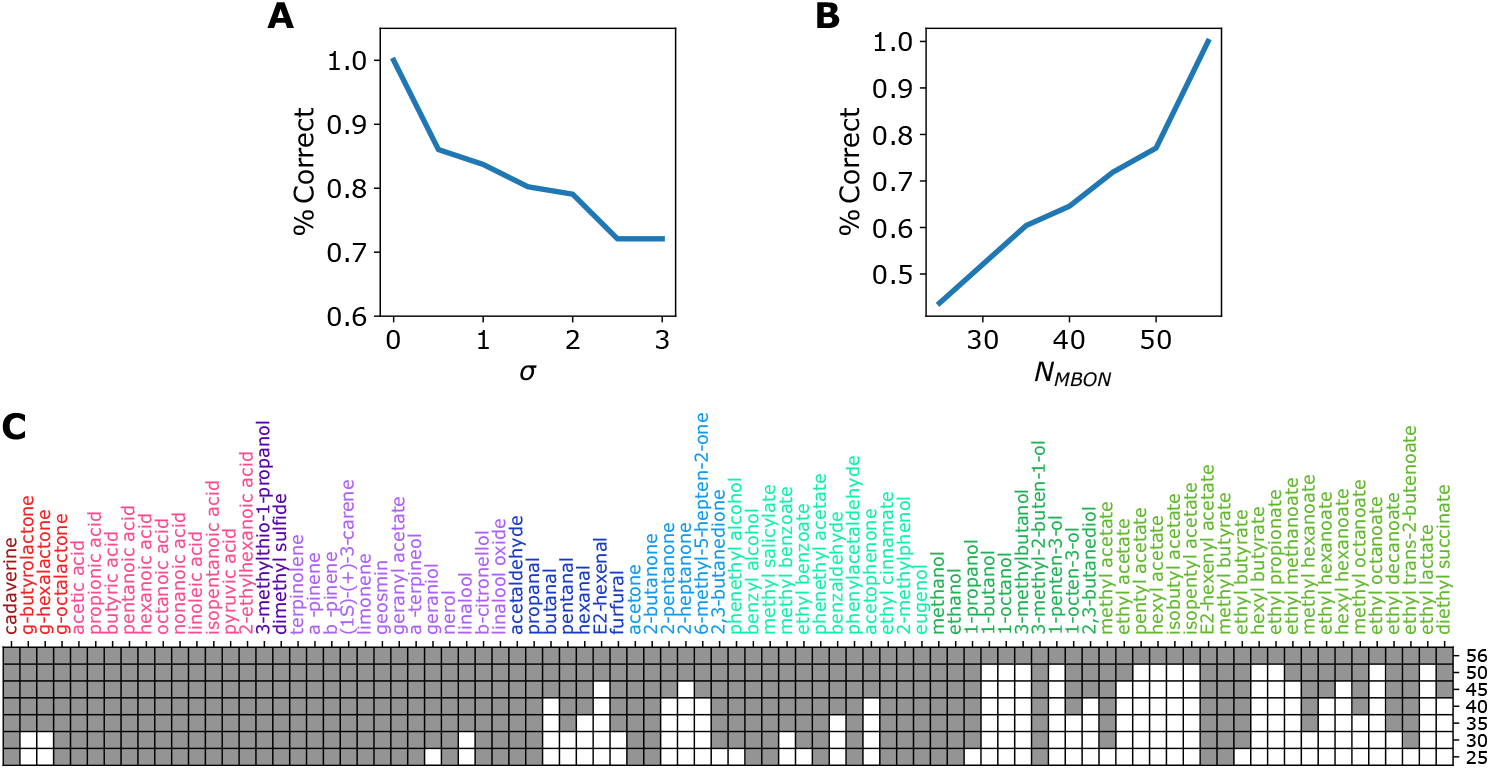
The robustness of odor identification. (A) Effect of noise on the success rate of odor identification. *σ* denotes the strength of noise added to the uPN activity profile. (B) Testing the odor identifiability using a random subset of MBONs of the size *N*_MBON_. The relationship between *N*_MBON_ and the success rate of odor identification. For a given odorant and a given size *N*_MBON_, MBONs are sampled ten times, where the odor identification is considered successful if at least one set of the sampled MBONs can accurately identify the given olfactory stimulus. We employ the 86 odorants where the identification was successful in the original setup. (C) The results of the identification test on 86 odorants from a subset of MBONs with the size *N*_MBON_ labeled on the *y*-axis. Gray denotes success and white denotes failure.

---

**TABLE S1.**
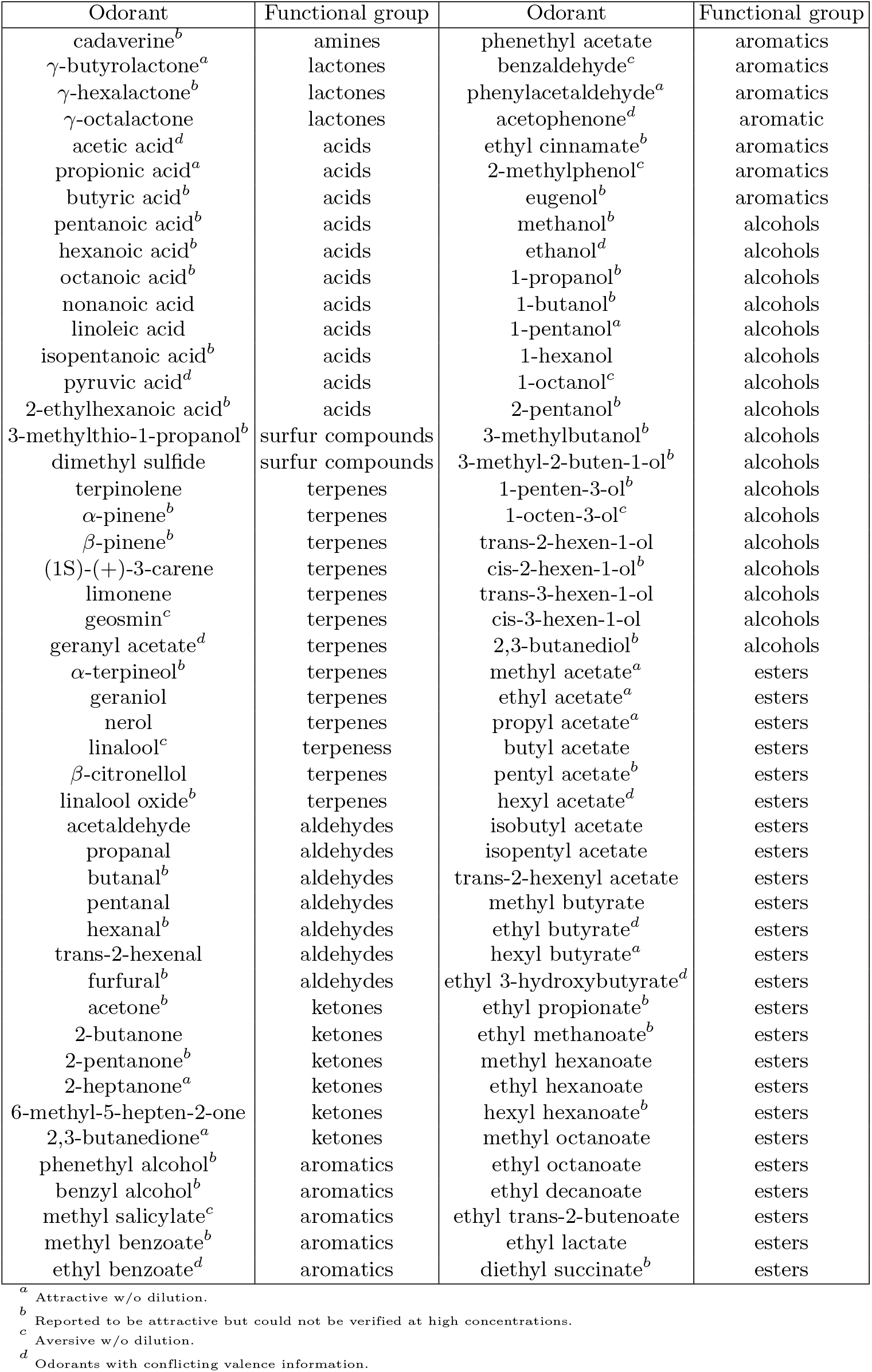
The list of odorants.

**Figure S1.**
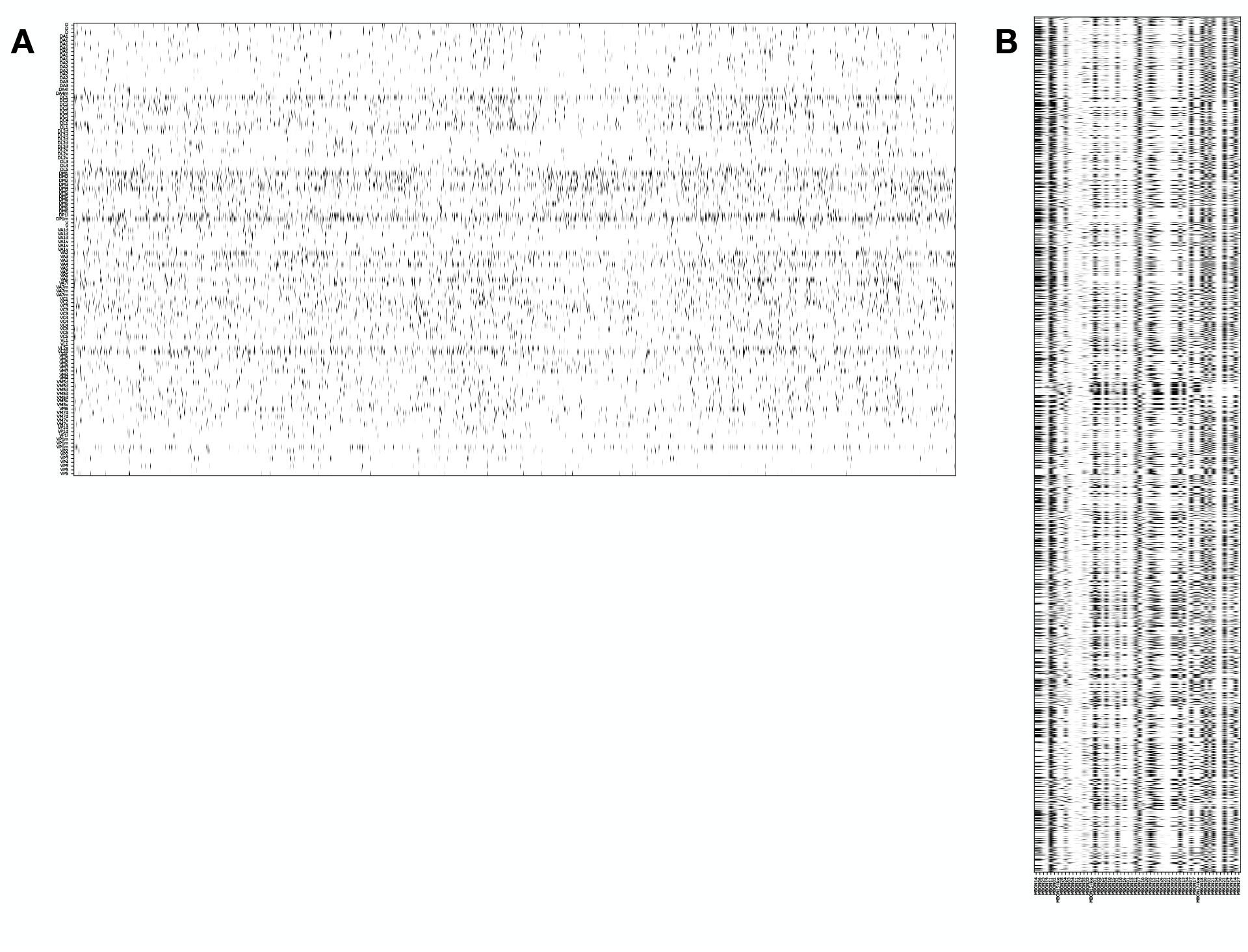
The synaptic connectivity matrices. (A) The 119 *×*1776 PN-KC connectivity matrix (𝒞^PN−KC^) and (B) the 1776 *×*56 KC-MBON connectivity matrix (𝒞^KC−MBON^). The values are normalized to the mean synaptic weight for better visibility.

**Figure S2.**
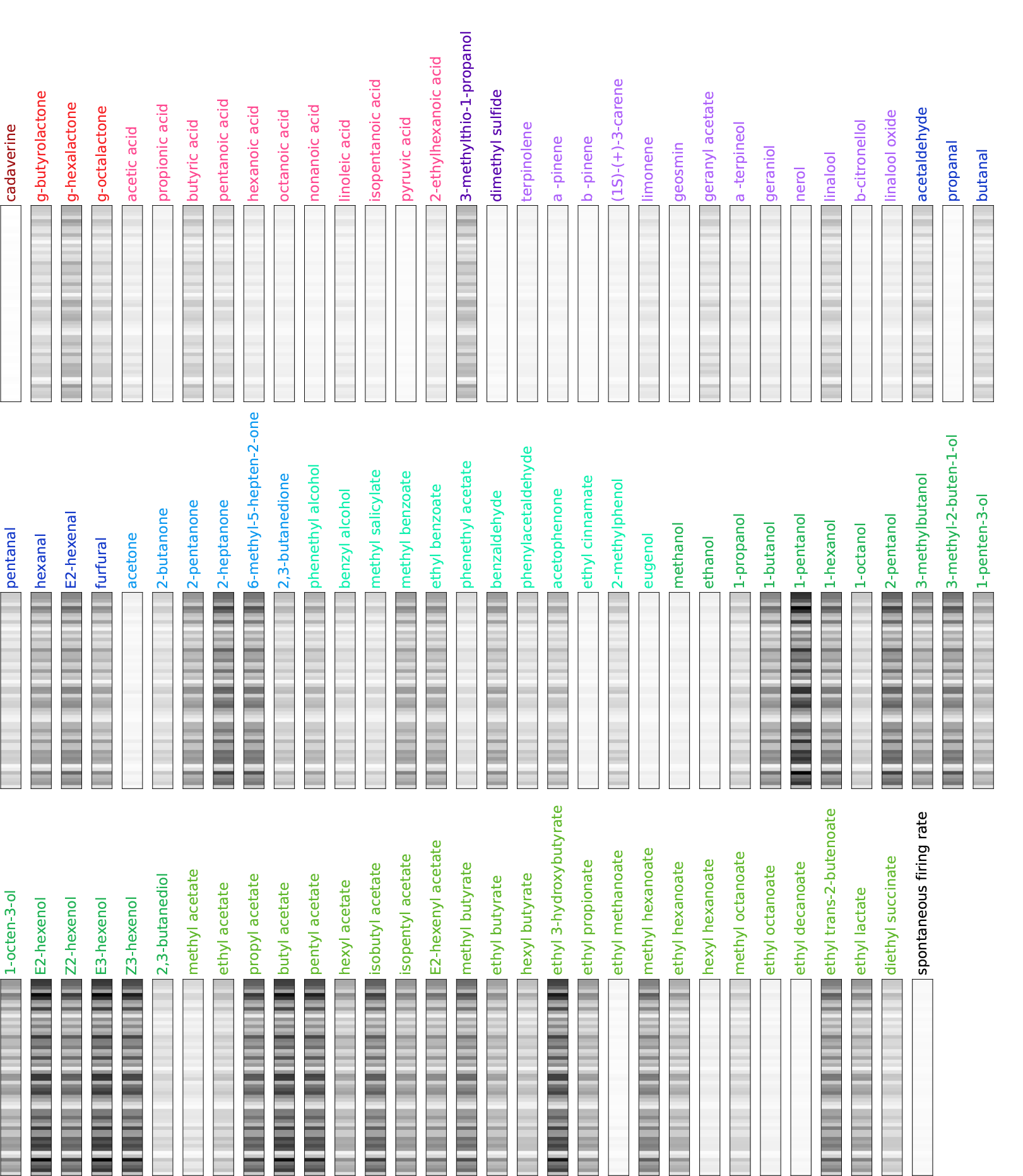
The MBON response profiles of all tested single odorants. The labels are color-coded based on the same odor categorization used by Hallem *et al*. [25] (Dark red: amines, red: lactones, pink: acids, purple: sulfur compounds, violet: terpenes, dark blue: aldehydes, blue: ketones, emerald: aromatics, green: alcohols, olive: esters)

**Figure S3.**
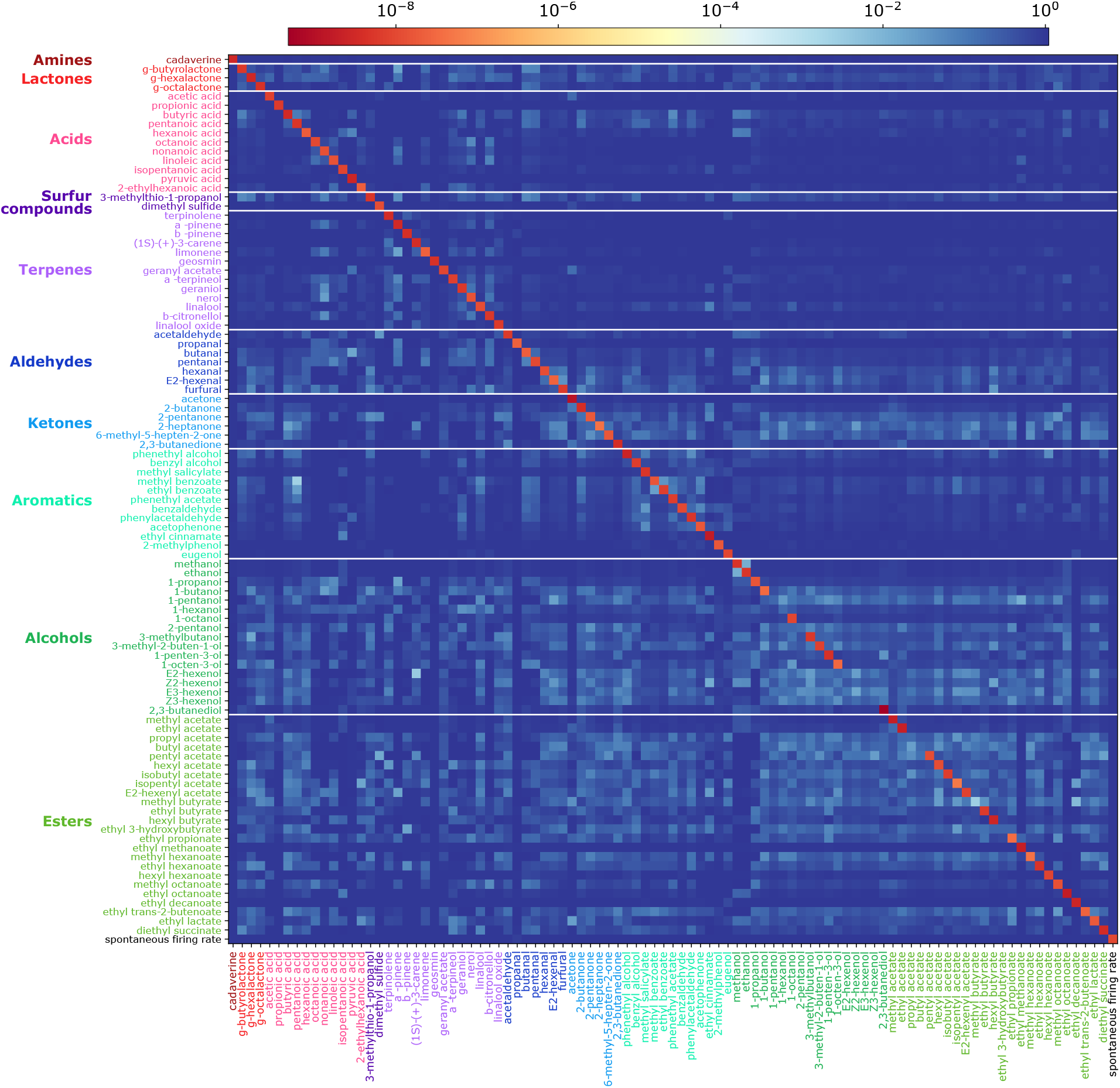
Each residual spectrum, calculated for the 96 odorants in Fig. 4 are represented along each row of the matrix. The residual *r*_*α*|*β*_ calculated for an input odorant *α* against other odorants *β* denoted in the horizontal axis. The same color-codes as Fig. 3 are used.

**Figure S4.**
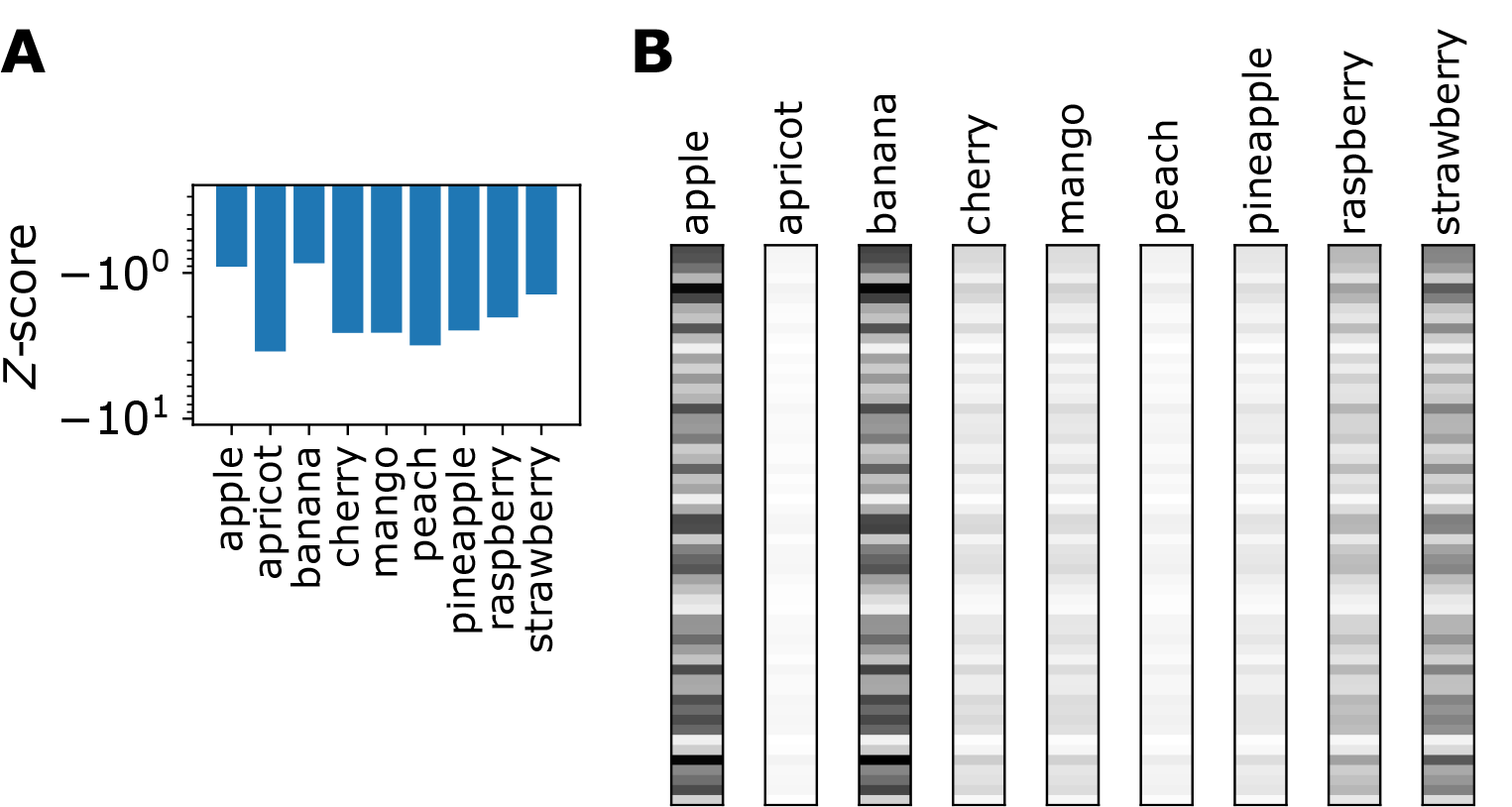
The output of the natural mixtures at 10^−2^ dilution. (A) The *Z*-scores. (B) The MBON response profiles.

**Figure S5.**
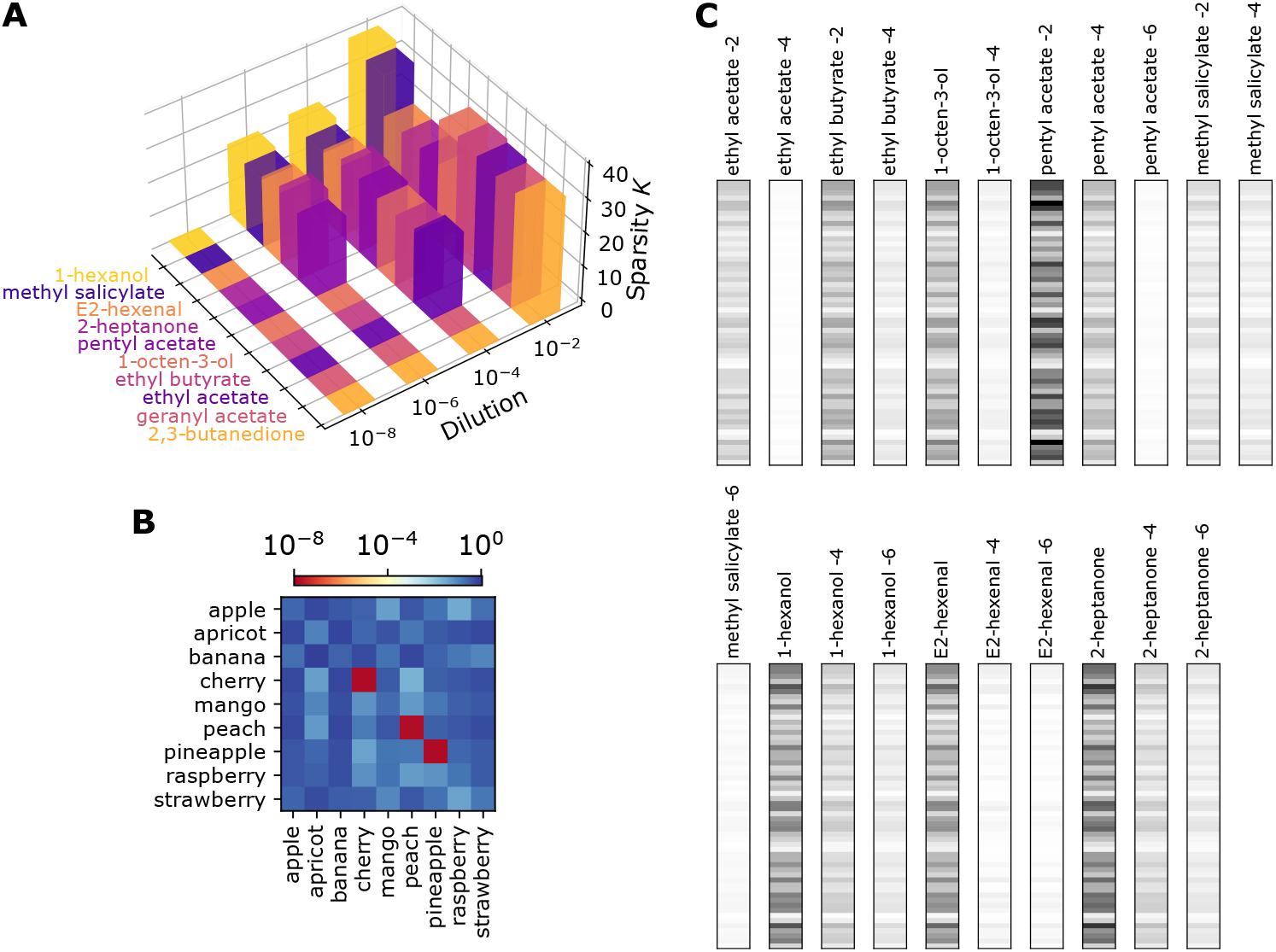
The effect of odorant concentration on odor perception. (A) The sparsity *K* for select odorants at varying dilutions. (B) The residuals *r*_*α*|*β*_ for undiluted fruit extracts. Each row depicts the spectrum of the residuals of an odor input *α* with respect to other test odors *β* (*β* = apple, apricot, …) given on the horizontal axis. (C) The MBON response profiles of single odorants at various dilution levels. −2, −4, and −6 signify the dilution levels of 10^−2^, 10^−4^, and 10^−6^, respectively.

